# Phenotypic and genomic signatures of interspecies cooperation and conflict in naturally-occurring isolates of a model plant symbiont

**DOI:** 10.1101/2021.07.19.452989

**Authors:** Rebecca T. Batstone, Liana T. Burghardt, Katy D. Heath

**Affiliations:** Carl R. Woese Institute for Genomic Biology, University of Illinois at Urbana-Champaign, 1206 West Gregory Drive, Urbana, IL 61801 USA; Department of Plant Science, The Pennsylvania State University, 103 Tyson Building, University Park, PA, 16802 USA; Department of Plant Biology, University of Illinois at Urbana-Champaign, 286 Morrill Hall, 505 South Goodwin Avenue, Urbana, IL 61801 USA

**Keywords:** conflict, cooperation, GWAS, mutualism, pleiotropy

## Abstract

Given the need to predict the outcomes of (co)evolution in host-associated microbiomes, whether microbial and host fitnesses tend to trade off, generating conflict, remains a pressing question. Examining the relationships between host and microbe fitness proxies at both the phenotypic and genomic levels can illuminate the mechanisms underlying interspecies cooperation and conflict. We examined naturally-occurring genetic variation in 191 strains of the model microbial symbiont, *Ensifer meliloti*, paired with each of two host *Medicago truncatula* genotypes in single- or multi-strain experiments to determine how multiple proxies of microbial and host fitness were related to one another and test key predictions about mutualism evolution at the genomic scale, while also addressing the challenge of measuring microbial fitness. We found little evidence for interspecies fitness conflict; loci tended to have concordant effects on both microbe and host fitnesses, even in environments with multiple co-occurring strains. Our results emphasize the importance of quantifying microbial relative fitness for understanding microbiome evolution and thus harnessing microbiomes to improve host fitness. Additionally, we find that mutualistic coevolution between hosts and microbes acts to maintain, rather than erode, genetic diversity, potentially explaining why variation in mutualism traits persists in nature.

---

Recent advances in sequencing and microbiome research have revealed the ubiquity of microbial symbioses, meaning that many important host phenotypes, such as plant yield in agriculture or disease-related traits in humans, are actually **symbiotic extended phenotypes**, their variation being influenced by loci present within interacting microbial symbionts in addition to the host (1–6). When loci influence fitness-related traits of both host and symbiont, which we henceforth refer to as **symbiotic pleiotropy**, they determine the degree to which partners’ fitnesses are aligned (i.e., same-sign or concordant effects) or in conflict (i.e., opposite-sign or discordant effects). Identifying the loci underlying symbiotic pleiotropy is therefore critical not only for illuminating the genetic basis of symbiotic extended phenotypes, but also for predicting how hosts and symbionts coevolve in nature.

In symbiotic mutualisms, wherein partners trade fitness benefits (7), whether fitness conflict or alignment drives the evolution of these interactions is hotly debated (8–13). Many models of mutualism rely on the key assumption that cooperation is inherently costly and thus that selection should favour less-cooperative, potentially ‘cheating’, partners that forgo paying the costs whilst continuing to receive benefits (14–17). At the phenotypic level, cheaters would be seen as symbiont genotypes that gain fitness at the host’s expense (i.e., points in the bottom right quadrant of **Fig. 1A**), while fitness conflict would be seen as an overall negative correlation (grey line in **Fig. 1A**). In contrast, cooperators would be seen as symbiont genotypes whose increase in fitness is associated with an increase in host fitness (i.e., points in the top right quadrant of **Fig. 1A**), while fitness alignment would be seen as an overall positive correlation (orange line in **Fig. 1A**). Evidence for fitness conflict within mutualism is mixed: although several studies have identified symbiont genotypes that gain fitness at their host’s expense (e.g., 18, 19), recent experimental evolution studies instead found that microbial adaptation to particular host genotypes is associated with an increase, rather than a decrease, in host fitness (20–22). Yet, fitness alignment at the phenotypic level does not necessarily preclude fitness conflict at the genomic level: rather than dichotomous categories of “cooperator” or “cheater”, mutualist genomes are best viewed as mosaics of loci (5), some underlying cooperation while others underlie conflict. Whether coevolution resulting from symbiosis leads to more beneficial interactions and greater mutualism stability, or alternatively, more antagonism and less stable interactions, therefore requires examining the relationships between host and symbiont fitness proxies at both the phenotypic and genomic levels.

**Fig. 1.**
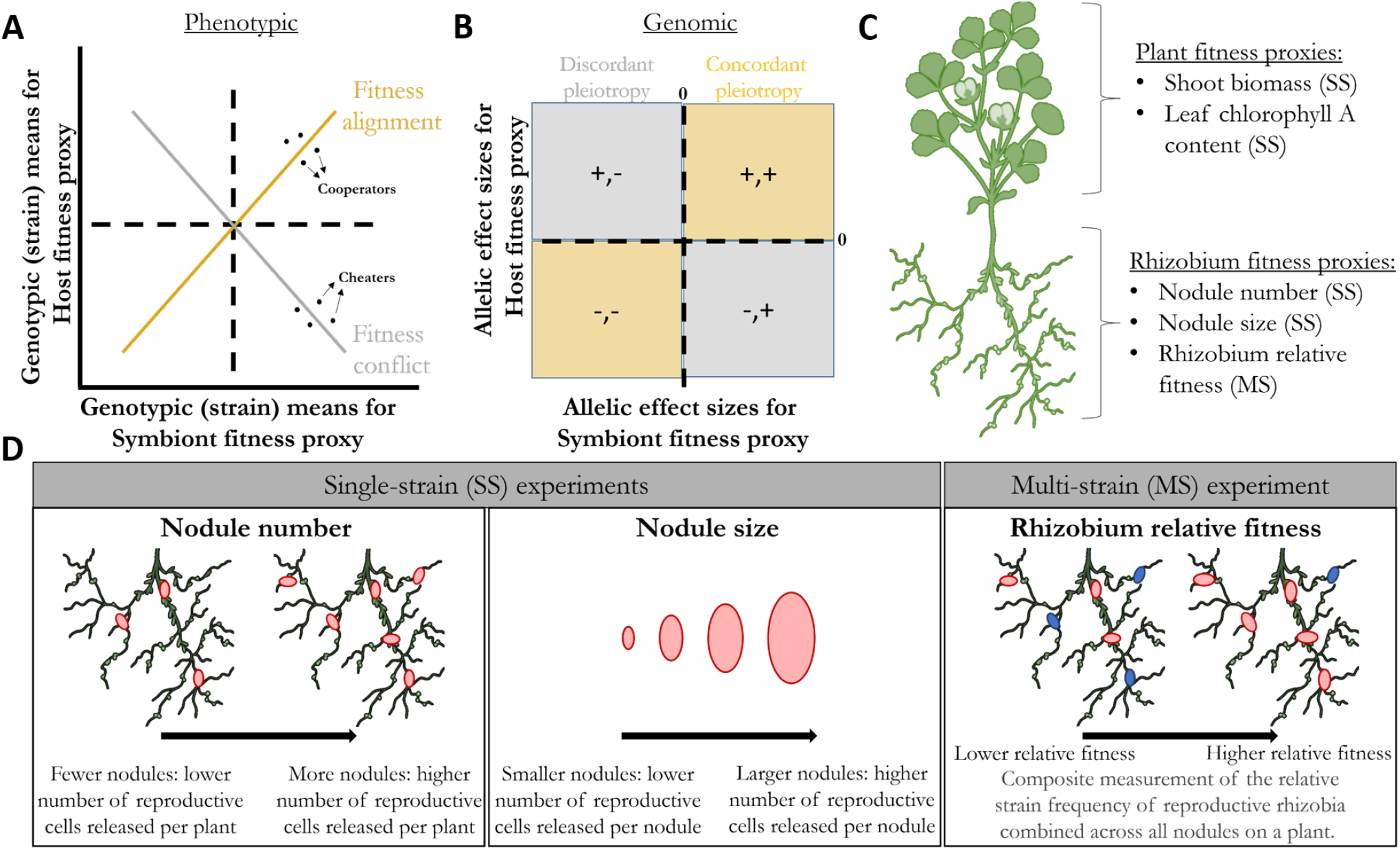
Interpreting patterns of fitness alignment and conflict at both the phenotypic and genomic levels. **A)** Phenotypic-level correlations of genotypic (strain) means for rhizobium fitness proxies (e.g., nodule number, nodule size, rhizobium relative fitness) or plant fitness proxies (shoot biomass or leaf chlorophyll A content) on the x- and y-axes, respectively. The positive (orange) or negative (grey) correlations represent fitness alignment or conflict, respectively. **B)** Genomic-level correlations of allelic effect sizes determined in GWAS for rhizobium fitness proxies or plant fitness proxies on the x- and y-axes, respectively. Points appearing in orange or grey quadrants represent pleiotropic variants with concordant (same-sign) or discordant (opposite sign) effects, respectively, on both host and symbiont. **C)** Plant and rhizobium fitness proxies measured, parentheses indicating the experiment type for which proxies were measured (SS = single-strain; MS = multi-strain). **D)** Rhizobium fitness proxies corresponding to the pink rhizobium strain. In single-strain experiments, nodule number indicates the number of reproductive cells released per plant, while nodule size indicates the number of reproductive cells released per nodule. In the multi-strain experiment, rhizobium relative fitness is a composite metric that combines competition among strains for nodule occupancy and for host resources once in nodules. Nodules pink versus blue in colour are used here to illustrate two different rhizobium strains competing, however, a total of 89 strains were inoculated together onto plants in the multi-strain experiment.

Genome-wide association studies (i.e., GWAS) can be used to reveal the genes, as well as specific segregating mutations (i.e., variants), that underlie variation in symbiotic extended phenotypes in natural populations (20, 23–27). Because they provide an estimate of both the strength and direction of effects of particular alleles on the trait, henceforth referred to as **allelic effect sizes**, GWAS are especially useful for identifying loci underlying symbiotic pleiotropy, and thus, defining the mutational landscape of mutualism evolution in nature. For example, if symbiotic pleiotropy is extensive and its effects on fitness-related traits in the interacting partners tend to be discordant (grey quadrants in **Fig. 1B**), then conflict should underlie the evolution of mutualism, allowing for the possibility of cheating individuals that are competitively superior, as mutualism theory predicts (14–17). In contrast, if pleiotropic effects are overwhelmingly concordant (orange quadrants in **Fig. 1B**), fitness alignment rather than conflict should be the null hypothesis in mutualism, and models relying on cheating genotypes as the main driver of mutualism evolution may not be suitable.

A longstanding mutualism paradox is that host-driven selection for the ‘best’ symbiont genotype should reduce overall symbiont diversity, yet diverse symbiont populations persist in nature (reviewed by 28, 29). Identifying patterns of selection acting on loci that determine fitness outcomes in natural populations could be key for resolving this paradox. Studies to date examining patterns of molecular variation have found stabilizing or purifying selection acting on candidate genes associated with partner recognition or quality, rather than patterns suggesting rapid turnover of alleles underlying conflict (30–33). In a recent in silico GWAS (5), conflict over mutualistic trait optima between hosts and microbes tended to increase genetic variance due to repeated sweeps, while alignment scenarios resulted in stabilizing selection and decreased genetic variance as trait optima were reached (5). Because GWAS has the power to reveal individual allelic effects, it can be used to identify loci underlying fundamental sources of conflict at the genomic level (e.g., grey quadrants, **Fig. 1B**) even if fitness alignment is realized at the phenotypic-level (e.g., orange line, **Fig. 1A**).

Over the past ~25 years, legume-rhizobium symbioses have been developed as models for understanding mutualism evolution (34–37). This interaction is one of the most ecologically and economically important symbioses, contributing upwards of 20 million tonnes of nitrogen (N) to the global N-cycle, and saving billions of dollars that would have otherwise been spent on synthetic N fertilizer production (38). Legumes house rhizobia within specialized root nodules and supply them with photosynthate, while rhizobia within the nodules convert atmospheric N to a plant-usable form, resulting in a beneficial exchange of resources. Key traits of this symbiosis, such as plant biomass and nodule number, are known to be influenced by variants in the genomes of both host and microbes, as well as the epistatic interactions between them (Genotype-by-Genotype or G x G interactions; e.g., 1, 26, 34, 39–42), making this symbiosis an excellent model for understanding mutualistic coevolution.

A quantitative comparison of single-strain and multi-strain proxies of rhizobium fitness, as well as their correlations with plant fitness, is needed to reveal interspecies conflict. Seed number, or close correlates such as aboveground (shoot) biomass and leaf chlorophyll A content, are well-established proxies for reproductive fitness in annual plants (23, 43, 44) (**Fig. 1C**). In contrast, estimating rhizobium fitness has been an empirical and conceptual challenge (8–10, 45, 46). Early attempts to estimate rhizobium fitness relied on single-strain experiments, whereby plants were inoculated with a single rhizobium strain, and fitness proxies including nodule number and nodule size were measured (**Fig. 1D**). Both measures reflect rhizobium fitness because a rhizobium that establishes a nodule in symbiosis can gain a fitness benefit on the order of 10^5^ – 10^7^ (47), larger nodules release more rhizobia (43, 46, 48), and intuitively, strains that produce more nodules will release more rhizobia (1, 37, 48). While rhizobium fitness proxies measured in single-strain experiments can be directly correlated with host benefit (e.g., shoot biomass), these measures have been criticized for producing spurious positive correlations due to stronger fitness feedbacks between host plants and rhizobia (9). In contrast, multi-strain experiments decouple individual strain fitness from host growth and better reflect rhizobia fitness in natural and agricultural soils where many strains coexist. Recent advances merging population genomics and metagenomics have enabled measuring **rhizobium relative fitness**, i.e., a strain’s ability to compete for nodulation opportunities and extract host resources once in nodules when other strains are present (25, 49). Multi-strain experiments that have estimated rhizobium relative fitness have so far hinted at a surprising lack of correlation between single-strain and multi-strain measures of rhizobium fitness (25), motivating our comprehensive analysis here.

Here we combine two datasets: first, GWAS that test for the association between rhizobium variants and both plant and rhizobium fitness proxies measured in single-strain experiments (**Fig. 1C**), whereby 191 strains of *Ensifer meliloti* collected from natural populations were inoculated individually onto one of two genotypes of the host plant *Medicago truncatula*. Second, a GWAS based on a new dataset that measured rhizobium relative fitness in a multi-strain experiment (**Fig. 1D**), whereby 89 of the 191 *E. meliloti* strains were inoculated together onto the same two host genotypes. We combine these datasets to first ask: What are the relationships among rhizobium fitness proxies measured in both single-strain (nodule number, nodule size) and multi-strain (rhizobium relative fitness) experiments? Are these proxies genetically distinct and thus likely to evolve independently, or are they linked through pleiotropy, and thus, evolve together? We use this information to next address the potential for genomic conflict in this symbiosis, asking: Do variants tend to have aligned or conflicting effects on host and symbiont fitnesses? Finally, we ask whether there is any evidence that historical selection has differentially shaped loci underlying fitness alignment versus conflict, as we might predict under different models of mutualism evolution.

## Materials and Methods

### Study system

Full details are provided in Riley et al. (50) and **SI Methods**. *Ensifer* (formerly *Sinorhizobium*) *meliloti* is an Alphaproteobacteria that forms symbiosis with various *Medicago* spp., fixing atmospheric N in return for plant photosynthate. All 191 strains used here were collected from the native range of the host *Medicago truncatula*, spanning Spain to France (as detailed in 50). *E. meliloti* has a multipartite genome, including a large (~3.7 Mbp) chromosome and two symbiosis plasmids (pSymA and pSymB, ~1.4 Mbp and ~1.7 Mbp, respectively); pSymA contains many of the canonical genes for nodulation and N-fixation (51, 52). We used two lines of *M. truncatula* DZA 315.16 (hereafter DZA) and A17 in separate single-strain experiments and a multi-strain experiment detailed below.

### Single-strain experiments

Full details are provided in Batstone et al. (42), and **SI Methods**. Briefly, we conducted two separate experiments in Sept and Nov 2018, one for each *M. truncatula* line, DZA and A17, respectively. At least three replicate seeds of each plant line were planted individually into pots (most treatments having four to five plant replicates) and were singly inoculated with one of 191 strains of *E. meliloti* described above. Within each experiment, we measured proxies for both plant fitness (shoot biomass, leaf chlorophyll A content) and rhizobium fitness (nodule number, per nodule weight; **Fig. 1C,D**).

### Multi-strain experiment

We grew A17 and DZA hosts with a multi-strain inoculum composed of 89 of the 191 strains used in the single-strain experiments (described in more detail in **SI Methods**). Using the “select-and-resequence” approach (25), this experiment allowed us to generate new data on **rhizobium relative fitness**, which represents a strain’s ability to both compete for nodulation opportunities and extract host resources once in nodules when 88 other strains were present (**Fig. 1D**). This fitness proxy was obtained by sequencing pooled nodule samples from each plant at the end of the experiment and estimating each strain’s frequency using a haplotype reconstruction method (53). We then obtained each strain’s relative fitness by calculating the fold change in the frequency of a strain at the end of the experiment relative to its mean frequency at the beginning (see **SI Methods**). Because we wanted our rhizobium relative fitness metric to represent which strains will be present in the soil in subsequent generations, our method focuses on measuring strain frequencies of undifferentiated rhizobia in nodule pools; while differentiated bacteroids are responsible for fixing N, they are reproductively sterile, and thus, do not contribute to the next generation. Although the plant and microbe growth conditions used here differed slightly from those used in the single-strain experiments (due to these experiments being conducted in separate places and times), previous work has shown that strain frequencies are stable in response to environmental variation within an experiment, between experiments, and across host generations (49, 54). Thus, environmental variation due to differences in growth conditions is unlikely to significantly influence how strains behave across experiments.

### Phenotypic analyses

As described in Batstone et al. (42), we calculated the estimated marginal means for each fitness proxy in each experiment (i.e., nodule number, nodule size, rhizobium relative fitness, shoot biomass, leaf chlorophyll A content), correcting for the effect of rack, using the emmeans package (v1.4.1, 55) in the R environment (56). We then conducted linear pairwise regressions (lm option in base R) for each fitness proxy against the other within each plant line.

### DNA isolation, whole-genome sequencing, and variant calling

Detailed methods are provided in Riley et al. (50), and **SI Methods**. We obtained DNA from each of the 191 rhizobium isolates, sequenced their full genomes including the chromosome and two symbiosis plasmids, used a common reference-based assembly pipeline to align sequences and call single nucleotide polymorphisms (SNPs), and filtered the resulting SNPs based on sequence quality, depth, minor allele frequency, and missingness, resulting in a total of 36,526 filtered SNPs.

### Genome-wide association tests

Detailed methods are provided in **SI Methods**. We conducted linear mixed models (LMMs) implemented in GEMMA (v0.98.1, 57) to quantify **allelic effect sizes** that represent both the strength and direction of the association between variants and fitness proxies after correcting for rhizobium population structure. We ran ten separate association analyses in GEMMA, one for each of the four fitness proxies measured in single-strain experiments (**Fig. 1C**), plus rhizobium relative fitness measured in the multi-strain experiment, and for both host plant lines (DZA, A17; five proxies x two hosts = ten runs).

### Genomic analyses

We first identified pleiotropic variants as those that were significantly associated with more than one trait on the same host line, significance determined using a permutation method (described in more detail in the **SI Methods**). Variants that were significantly associated with two or more rhizobium fitness proxies were categorized as underlying **rhizobium fitness pleiotropy**, whereas variants associated with at least one rhizobium AND plant fitness proxy were categorized as underlying **symbiotic pleiotropy**. We further categorized whether pleiotropic variants had discordant (opposite-sign; +,- or -,+) or concordant (same-sign; +,+ or -,-) effects on pairwise fitness proxy combinations (**Fig. 1B**). Finally, to test whether selection acted on genes containing variants associated with rhizobium fitness and symbiotic pleiotropy, we used the R package PopGenome (v2.7.5, 58) to compute several commonly used test statistics that can detect signatures of historical selection and/or departures of neutrality, namely, nucleotide diversity (i.e., *π*), Tajima’s D, as well as Fu and Li’s F and D (59, 60). Additional details appear in **SI Methods**.

### Data availability

Strains and plant lines are available upon request. All raw data and analysis code are available on GitHub (see “Genetics_conflict_cooperation” folder). The Supplementary Information doc contains additional methods, results, figures and tables, as well as descriptions of datasets that have been uploaded along with this manuscript, and can be additionally accessed from bioRxiv and GitHub. Once raw sequence reads and assemblies are archived and made accessible on NCBI, accession numbers will be added to this manuscript.

## Results

### Relationships among rhizobium fitness proxies

In single-strain experiments, when we regressed strain means for nodule number and nodule weight, we found significant negative correlations for both hosts (**Fig. 2A**, left), indicating that strains creating larger nodules tended to form fewer total nodules on both host genotypes. At the genomic-level, most variants had discordant effects (**Fig. 2A**, right), similarly indicating a trade-off whereby variants that were positively associated with nodule weight tended to be negatively associated with nodule number, or vice-versa.

**Fig. 2.**
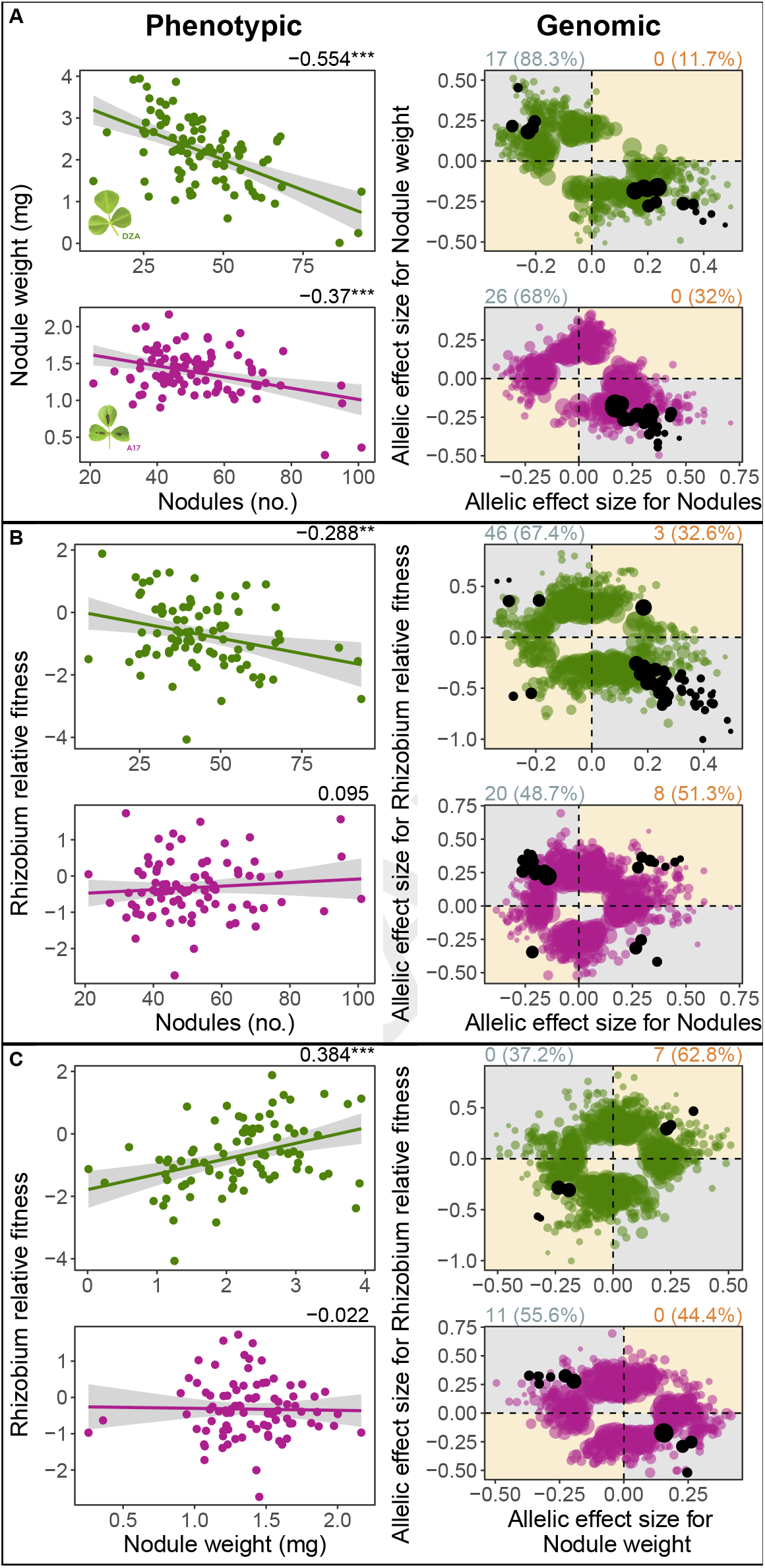
Trade-offs among rhizobia (*Ensifer meliloti*) fitness proxies prevail at both the phenotypic (left panels) and genomic-levels (right panels). **Phenotypic**: Genetic correlations between pairwise fitness proxies measured on plant lines DZA (green, top rows) or A17 (pink, bottom rows), based on 89 *Ensifer meliloti* strains. Dots represent estimated marginal strain means for nodule number and nodule weight, both being measured in single-strain experiments, or medians for rhizobium relative fitness measured in multi-strain experiments. Numbers at top right of each correlation represent Pearson correlation coefficients, while asterisks represent significance: * = p < 0.05; ** = p < 0.01; *** = p < 0.001. **Genomic**: Dots represent allelic effect sizes (i.e., beta scores calculated in GEMMA), those falling along the diagonal (orange quadrants) or off-diagonal (grey quadrants) represent variants with concordant or discordant effects, respectively. Coloured dots represent variants that were significantly associated with one of the two fitness proxies, while black dots represent pleiotropic variants, i.e., significantly associated with both fitness proxies. Numbers outside of and percentages within parentheses at the top left and right of each plot represent the pleiotropic variant counts and the proportion of total significant variants, respectively, that are discordant (left, in grey) or concordant (right, in orange).

Comparing single- and multi-strain experiments, when we regressed strain means for rhizobium relative fitness and nodule number, we found a weak but significant negative relationship for host line DZA only (**Fig. 2B**, left), indicating that strains that were more common in the multi-strain experiment formed fewer nodules in the single-strain experiment. At the genomic-level on DZA, most variants had discordant effects (**Fig. 2B**, right), again indicating a trade-off between nodule number and rhizobium relative fitness. For host line A17, most pleiotropic variants (20/28) had discordant effects (**Fig. 2B**, right), despite no relationship between these fitness proxies at the phenotypic level (**Fig. 2B**, left).

We found a significant, positive relationship between strain means for rhizobium relative fitness and nodule weight, again for DZA only (**Fig. 2C**, left). This result indicates that strains that were more commonly found in nodules in the multi-strain experiment formed larger nodules in the single-strain experiment. At the genomic-level, all pleiotropic variants were concordant (i.e., appearing in the top right or bottom left quadrants of **Fig. 2C**, right). For A17, while we found a lack of significant phenotypic correlation between nodule weight and rhizobium relative fitness (**Fig. 2C**, left), at the genomic level, all pleiotropic variants had discordant effects on these fitness proxies (**Fig. 2C**, right), indicating a trade-off at this scale.

These results overall support trade-offs at the phenotypic level, plus underlying discordant pleiotropy at the genomic level, suggesting that strains for which plants form numerous small nodules in single-strain experiments are less able to proliferate within and compete for nodules in multi-strain experiments. We additionally found host-dependent relationships among rhizobium fitness proxies, especially at the genomic level when regressing rhizobium relative fitness and nodule weight (**Fig. 2C**, right). However, we found fewer total pleiotropic variants associated with this relationship compared to the other two fitness proxy combinations (**SI Fig. S2A**, right), suggesting that nodule weight and rhizobium relative fitness are largely governed by different molecular mechanisms, and thus, likely to evolve independently.

### Relationships between plant and rhizobium fitness proxies

We found no correlation, in either host, between strain means for nodule number and shoot biomass (**Fig. 3A** left), or nodule number and leaf chlorophyll A content on A17 (**SI Fig. S3A**, left). However we found a significantly negative correlation between nodule number and chlorophyll A for DZA (**SI Fig. 3A**, left). Assuming chlorophyll A is related to the N-fixation efficiency of each strain (61–63), this result suggests that strains forming more numerous (and smaller) nodules tended to fix less N. At the genomic level, most variants had concordant effects for nodule number and shoot biomass on DZA, whereas the opposite was true for the same proxy pair on A17 (**Fig. 3A**, right). Most variants had discordant effects on nodule number and chlorophyll A on both hosts (**SI Fig. S3A**, right).

**Fig. 3.**
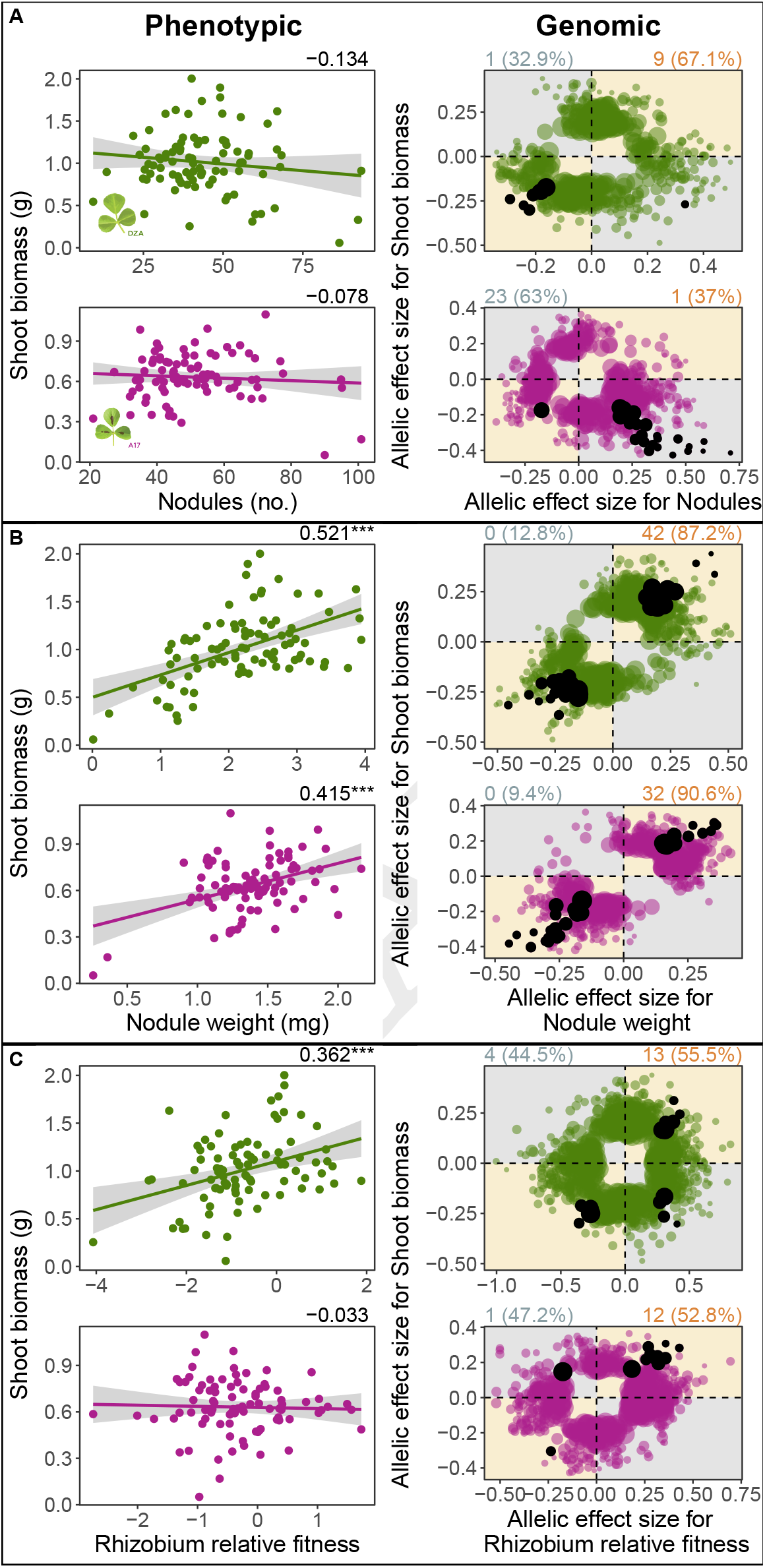
Fitness alignment between rhizobia (*Ensifer meliloti*) and host (*Medicago truncatula*) prevails at both the phenotypic (left panels) and genomic (right panels) levels. **Phenotypic**: Genetic correlations between pairwise fitness proxies measured on plant lines DZA (green, top rows) or A17 (pink, bottom rows), based on 89 *Ensifer meliloti* strains. Dots represent estimated marginal strain means for shoot biomass, nodule number, and nodule weight, all being measured in single-strain experiments, or medians for rhizobium relative fitness measured in multi-strain experiments. Numbers at top right of each correlation represent Pearson correlation coefficients, while asterisks represent significance: * = P < 0.05; ** = P < 0.01; *** = P < 0.001. **Genomic**: Dots represent allelic effect sizes (i.e., beta scores calculated in GEMMA), those falling along the diagonal (orange quadrants) or off-diagonal (grey quadrants) represent variants with concordant or discordant effects, respectively. Coloured dots represent variants that were significantly associated with one of the two fitness proxies, while black dots represent pleiotropic variants, i.e., significantly associated with both fitness proxies. Numbers outside of and percentages within parentheses at the top left and right of each plot represent the pleiotropic variant counts and the proportion of total significant variants, respectively, that are discordant (left, in grey) or concordant (right, in orange).

Strain means for nodule weight and shoot biomass were significantly positively correlated for both hosts (**Fig. 3B**, left). Nodule weight and chlorophyll A were significantly positively correlated on DZA, but uncorrelated on A17 (**SI Fig. S3B**, left). At the genomic level for both hosts, all pleiotropic variants had concordant effects for nodule weight and shoot biomass (**Fig. 3B**, right), as well as nodule weight and chlorophyll A (**SI Fig. S3B**, right).

Finally, rhizobium relative fitness and shoot biomass were significantly positively correlated for DZA but not for A17 (**Fig. 3C**, left), whereas rhizobium relative fitness and leaf chlorophyll A content were significantly positively correlated on both hosts (**SI Fig. S3C**, left). At the genomic level, most pleiotropic variants had concordant effects on rhizobium relative fitness and shoot biomass for both hosts (**Fig. 3C**, right), and this pattern was even stronger between rhizobium relative fitness and chlorophyll A on DZA with all but one pleiotropic variant having concordant effects (**SI Fig. S3C** right). Variant effects were mixed on A17 for rhizobium relative fitness and leaf chlorophyll A content (**SI Fig. S3C**, right).

Overall, we found mostly concordant relationships between rhizobium and plant fitnesses at both the phenotypic and genomic levels, suggesting a strong signal of fitness alignment in natural rhizobium populations. However, we note a lack of pleiotropic variants underlying the relationship between leaf chlorophyll A content and both rhizobium fitness proxies measured in single-strain experiments (i.e., nodule number and nodule weight; **SI Fig. S2C**, left & middle), suggesting these proxies are governed by different molecular mechanisms, and thus, should evolve independently.

### Selection acts differently on genes associated with alignment versus conflict

Using multiple diversity and neutrality metrics, we found that rhizobium genes associated with fitness alignment exhibit higher nucleotide diversity and stronger signatures of balancing selection compared to any other gene category analyzed. For three of the four test statistics, genes associated with concordant symbiotic pleiotropy (i.e., solid orange lines in **Fig. 4**) had significantly elevated values relative to the “null” (i.e., distributions in **Fig. 4**, all genes containing significant variants identified by GWAS). We did not see any significant deviations from the null for genes associated with discordant symbiotic pleiotropy (i.e., dotted grey lines in **Fig. 4**), or for genes associated with both concordant and discordant rhizobium fitness pleiotropy (**SI Table S1**, **SI Fig. S4**, **SI Dataset S1**).

**Fig. 4.**
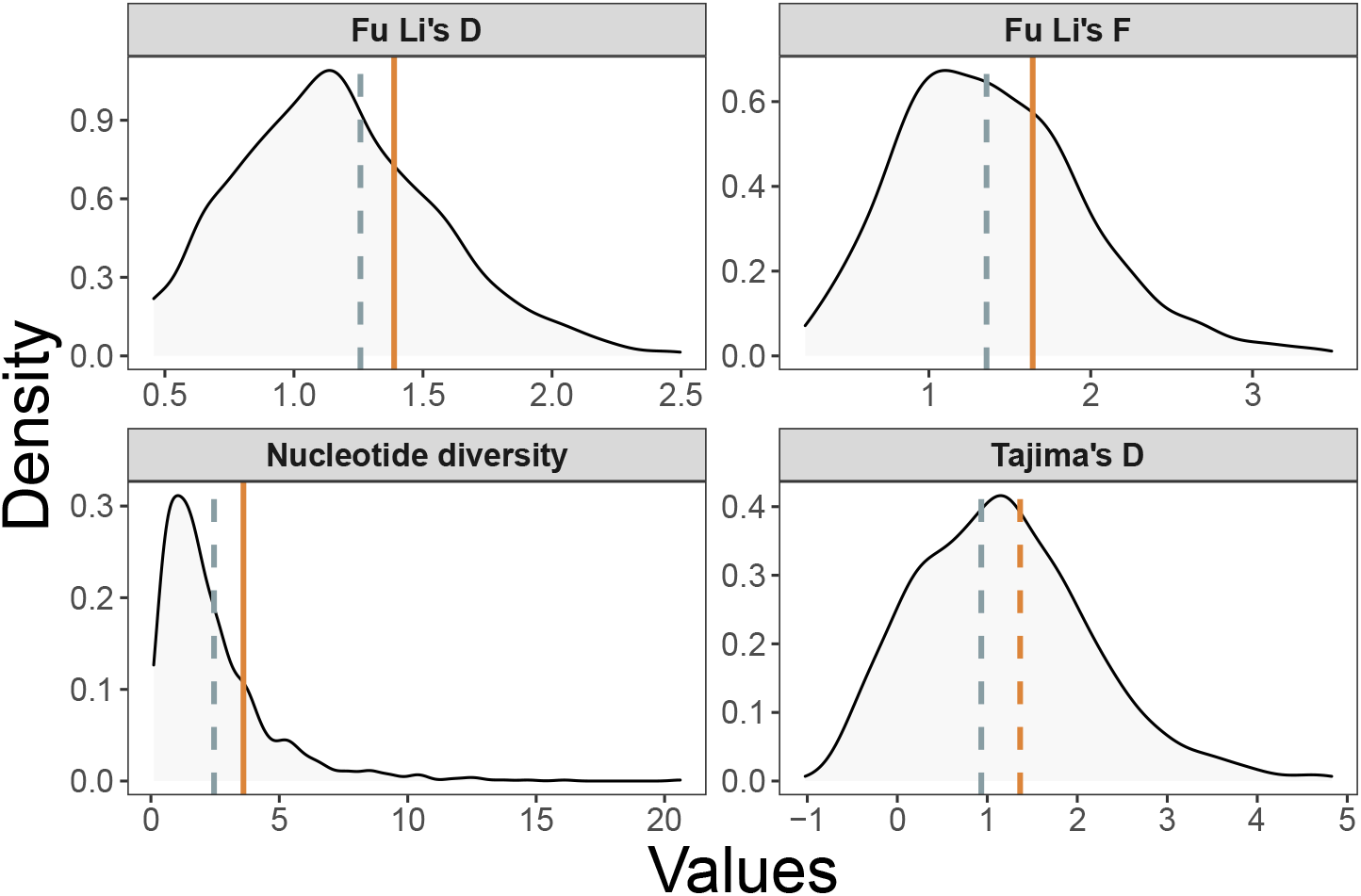
Neutrality statistics show elevated values for genes associated with fitness alignment. Vertical lines represent the average values calculated for four separate statistics, grey for discordant and orange for concordant genes. Distributions represent the same statistics calculated for all genes containing significant variants based on GWAS. Dashed and solid lines represent non-significant (p > 0.1) and significant (p < 0.1) differences, respectively, between each focal gene category and all significant genes (i.e., distributions).

### Loci associated with symbiotic pleiotropy are host-dependent

Comparing the identities and putative functions of variants associated with symbiotic pleiotropy on both host lines revealed little overlap – concordant variants giving rise to both high host and symbiont fitness (i.e., associated with fitness alignment) largely differed between the two host genotypes. Specifically, we identified a total of 168 variants associated with symbiotic pleiotropy, corresponding to 128 coding-genes (see **SI Dataset S2** for variant-level and **SI Table S2 & Dataset S3** for gene-level summaries). 60 and 93 of these variants were uniquely associated with fitness proxies measured on plant lines A17 and DZA, respectively, while only 15 variants were shared between hosts. We highlight some of the noteworthy genes identified in our analysis for each pleiotropic category in **SI Results**.

## Discussion

Leveraging genomics to quantify the genetic architecture underlying symbiotic extended phenotypes gives us the power to address long-standing issues in mutualism evolution with genome-scale resolution. Our results overall suggest that: 1) fitness alignment between hosts and rhizobia is common at both the phenotypic and genomic levels, with genes associated with alignment showing elevated nucleotide diversity and signatures of balancing selection; and 2) the lack of a relationship or even trade-offs between rhizobium nodule number and rhizobium relative fitness mean that measures of rhizobium fitness in multi-strain experiments should be prioritized when we want to predict rhizobium evolution. We discuss these main points in turn below.

### Alignment of host and symbiont fitnesses

In one-to-many symbioses, such as a single legume associating with a diverse population of rhizobia, less-beneficial symbionts are predicted to achieve higher relative fitness compared to more beneficial counterparts (14–16). While we find evidence for less beneficial rhizobia (i.e., points closer to zero along the y-axes of panels in **Fig. 3**), we find little evidence for “cheating” rhizobia genotypes or loci associated with an increase fitness at the host’s expense (i.e., a lack of points in the bottom-right quadrants of panels in **Fig. 3**). Instead, we found strong fitness alignment even in environments where multiple strains occupy the same plant.

Fitness alignment is ultimately governed by the degree to which mutualistic trait optima are shared among partners (5), as well as the degree of fitness feedbacks that enforce alignment between partners (11, 14–17). For example, legume host plants have autoregulation of nodulation to limit the formation of costly nodules once sufficient N levels are achieved (64). No such constraint exists for rhizobia; every nodule formed is expected to lead to greater potential fitness benefits. Thus, a mismatch between the optimum nodule number for a plant versus rhizobium could generate conflict (5, 65). Indeed, the strongest evidence for conflict in our and other studies (e.g., 1, 44, 48) comes from regressing plant fitness proxies on nodule number, suggesting conflict over a host’s total investment in nodulation.

In addition to controlling the total number of nodules formed, legumes can also allocate more carbon to nodules that fix more nitrogen (35, 66, 67), which acts to couple rhizobium quality and fitness even when multiple strains are present. Such fitness coupling mechanisms can be disrupted by mismatches between legume and rhizobium genotypes, allowing strains that fix little to no N to proliferate within nodules (19, 68, 69). Our observations of abundant alignment between host fitness proxies, nodule size, and rhizobium relative fitness suggest that such mismatches may be rare in nature, although in our study, neither host line was collected from the same sites as the rhizobia strains, and thus, are unlikely to naturally co-occur. Overall, our results suggest that trait optima can be shared even in one-to-many interactions, and that fitness feedbacks operating to align host and symbiont fitness are present even in diverse communities irrespective of coevolutionary history.

### Genomic resolution of conflict and alignment in symbiosis

Rather than to identify causal loci underlying symbiotic pleiotropy, the goal of our study was to examine broader patterns of fitness alignment or conflict across the genomes of numerous naturally occurring rhizobium strains. We found abundant concordant pleiotropic variants associated with both host and symbiont fitnesses, alongside evidence that selection has acted to maintain genetic variation within rhizobium genomes through time and/or space. A lack of discordant pleiotropic variants, like we have found, may have resulted from physiological constraints that make such variants impossible (i.e., alleles associated with larger nodules and small host biomass are rare or non-existent), or because of correlational selection that removes discordant variants from the population altogether (70, 71). In addition to direction, the extent of symbiotic pleiotropy (i.e., the number of pleiotropic variants) can inform whether traits are likely to evolve together or independently. Fitness proxy pairs with abundant pleiotropy (e.g., nodule size vs. shoot biomass) suggest a highly polygenic basis governed by many small-effect pleiotropic variants; other proxies with little pleiotropy (e.g., nodule number vs. chlorophyll) are largely governed by different sets of genetic mechanisms, and are thus likely to evolve independently.

One of our more critical findings, that the loci associated with both high host and symbiont fitness benefits (i.e., fitness alignment) largely differed across host genotypes, provides one solution to the mutualism paradox: if host genotypes act as distinct selective environments for rhizobia, meaning that the “best” symbiont genotype differs among host genotypes, then symbiont diversity could be maintained in the face of host selection. Host-dependent loci underlying fitness alignment were also found in experimentally evolved rhizobia isolates (20) and host-mediated balancing selection was previously proposed as a mechanism maintaining rhizobial diversity in native populations of *Bradyrhizobium* (72). Our results provide a solution to the mutualism paradox: symbiont diversity can be maintained via balancing selection acting on host-dependent rhizobium loci underlying fitness alignment.

We nonetheless found several instances of fitness conflict at the genomic level despite alignment at the phenotypic level (e.g., shoot biomass vs rhizobium relative fitness on DZA). Linkage disequilibrium (LD) could lead to discordant alleles being “packaged” into multilocus genotypes that show fitness alignment. Such LD results from multiple, non-mutually exclusive factors including epistatic interactions among individual variants that render discordant variants effectively neutral, and/or past selection favouring allelic combinations that increase both host and symbiont fitness (and disfavours discordant combinations; 73, 74).

Overall, these findings highlight the polygenic nature of symbiotic extended phenotype variation in nature — where the collective action of individual mutations, their additive and nonadditive effects (42), and a history of selection shapes the trait variation currently present in natural populations (75, 76). However, because of the limitations inherent within GWAS, including linkage disequilibrium that makes it difficult to pinpoint causal variants as well as false positive and negative associations, we acknowledge that the function of any specific locus identified here needs to be further validated in follow up experiments. Additionally, we only focus on allelic substitutions rather than presence/absence variation generated via gene gain and loss, the latter of which previous studies have found to be associated with exploitative traits in rhizobia (e.g., 68).

### Trade-offs among rhizobium fitness proxies and the rhizobium competition problem

Understanding how microbial symbioses, which are ubiquitous and important in nature, evolve (or coevolve) requires accurate estimates of symbiont fitness in ecologically realistic settings. Given our evidence at both the phenotypic and genomic levels for trade-offs among rhizobium fitness proxies, and because diverse strains of rhizobia co-occur in nature, relative fitness proxies should be used whenever possible (25). Nevertheless these proxies are not replacements for those measured in single-strain experiments because they cannot be used to assign individual genetic means for whole-plant measures of host benefit (plant biomass and seed number) to individual strains (e.g., 1, 26, 39, 42), necessitating that host benefit and rhizobium fitness be measured on separate individuals.

Together our results suggest that the genetic architectures associated with rhizobium fitness proxies, and their relationships, are host-dependent, and thus that their evolutionary trajectories are influenced not only by variation in the rhizobium’s genome, but also by variation in host traits. For example, host genotypes could differ in their ability to exert sanctions or partner choice, quantitative traits known to vary in host populations (44, 77). At the genomic-level, distinct variants underlying rhizobium fitness pleiotropy on each host genotype suggests that the genetic mechanisms (i.e., genes, pathways, metabolic processes) governing the relationship between fitness proxies are largely non-overlapping when rhizobia interact with different hosts. Such host genotype-dependent shifts in the rhizobium genetic architecture of these symbiotic extended phenotypes are supported by transcriptomics studies of G x G interactions (25, 40) and GWAS revealing distinct sets of candidate genes in different host backgrounds (26, 42). Similar genetic variation exists in hosts (78) and undoubtedly interacts with the variants we identify here, thus, should be accounted for if we want to uncover the multi-genomic basis of symbiotic extended phenotypes.

## ACKNOWLEDGMENTS

For their insightful feedback on the manuscript, we thank two anonymous reviewers and an associate editor. For greenhouse help, we thank Alex Riley, Hanna Lindgren, Cassandra Allsup, Laura Goralka, Mannix Burns, Brendan Epstein, and Amy-Marshall Colon, as well as the greenhouse staff, particularly Debbie Black. For feedback on the results and analyses, we thank Peter Tiffin, Brendan Epstein, and Robert Williamson. We acknowledge funding from the National Science Foundation (K.D. Heath: IOS-1645875 and NPGI-1401864; L.T. Burghardt: IOS-1724993 and IOS-1856744), a Carl R. Woese Institute for Genomic Biology postdoctoral fellowship to R.T. Batstone, and strain sequencing by Joint Genome Institute (CSP-1223795). This work was additionally supported by the USDA National Institute of Food and Agriculture Federal Appropriations under Project PEN04760 and Accession no. 1025611.

