## Supplementary Materials for "Phenotypic and genomic signatures of interspecies cooperation and conflict in naturally-occurring isolates of a model plant symbiont"

##### This PDF file includes:

Supplementary text
Legends for Dataset S1 to S3
Figs. S1 to S4
Tables S1 to S2
SI References

##### Other supplementary materials for this manuscript include the following:

Datasets S1 to S3

### Supporting Information Text

#### SI Methods

**Study system.** Host lines DZA and A17 were originally collected from Algeria and Australia, respectively, and thus, were not collected from the same sites for which our 191 rhizobium strains were isolated. However, these two host genotypes are well-characterized from previous studies investigating the role of nodule-specific cysteine rich (NCR) peptides in N-fixation, whereby strain-specific responses depend on each line's NCR genetic composition (1–4). These lines were also included in the *M. truncatula* HapMap (HM) panel (DZA corresponding to accession HM005, A17 to HM101), and are found in distinct clades within *M. truncatula*'s phylogeny (5).

**Single-strain experiments.** Seeds were prepared according to published protocols (6). Briefly, seeds were nick- or sandpaper-scarified, surface sterilized in EtOH for 30s, bleached (6% hypochlorite) for 7 min, rinsed with d$H_2O$ until bleach was completely removed, and imbibed in d$H_2O$ overnight in the dark at 4°C. For planting, we used SC10R Cone-tainers (Stuewe & Sons Inc., Tangent, OR), filling each with autoclaved potting media consisting of one part root wash (1:1:1 mix of field soil:torpedo sand:Turface) and one part Turface MVP calcined clay (Profile Products LLC, Buffalo Grove, IL, USA). A single seed was sown into each pot, and pots were completely randomized with respect to treatment across three greenhouse benches located at the University of Illinois at Urbana-Champaign (UIUC). We cultured rhizobia by loop-inoculating sterile liquid tryptone-yeast (TY) medium (7) with each strain individually, placing them into a shaking incubator set to 30°C and 200 rpm overnight. We measured the $OD_{600}$ of each culture, and diluted to $\sim 10^6$ cells per ml using sterile TY media to ensure all plants were inoculated with the same initial starting densities. Plants were misted throughout the experiment using upright misting nozzles (1/2" M NPT, Senninger Irrigation Inc., Clermont FL), and harvested six to seven weeks post-inoculation.

Within each experiment, we measured proxies for both plant (shoot biomass, leaf chlorophyll A content) and rhizobium (nodule number, per nodule weight) fitness. We measured chlorophyll using a SPAD 502 Plus (Spectrum Technologies, Inc. Aurora, IL, USA), recording the mean of three separate measurements on the more recently emerged leaf. Once plants were harvested, all nodules were counted, and ten nodules were haphazardly chosen to be weighed to the nearest 0.001 mg, dividing this number by ten to obtain a per-nodule average for each plant. Shoots were placed in a drying oven set to 60°C and left for three days, being weighed to the nearest 0.01 g. Contamination was minimal in either experiment: we only found 24 nodules on six internal controls (i.e., plants embedded within the experiment; range: 1-9 per plant), and no nodules on external controls (i.e., plants located on a bench adjacent to the experimental plants).

**Multi-strain experiment.** The 89 strains used in this experiment were chosen because they captured the maximum amount of rhizobial genetic diversity, providing the "select-and-resequence" (described more below, and in detail in 8) approach with strong discriminatory power; closely related strains are more challenging to differentiate. Within-plant strain diversity of nodules is typically high in nature, often greater than among plant or site diversity, and for the papers that have actually quantified strain diversity (9–12), 89 is a realistic number of strains to find in association with a plant.

We inoculated each strain into one of 89 culture tubes filled with 3 ml of TY media (6 g tryptone, 3 g yeast extract, 0.38 g $CaCl_2$ per L) (7) and grew cultures to stationary phase by placing the tubes in a 120 rpm shaker for 72 hours at 28°C. We combined all cultures, mixed well, and used this mixture to inoculate 3-day old seedlings with $\sim 10^7$ rhizobial cells diluted in 25 ml of N-free fertilizer. Three days prior to inoculation, we filled twelve Cone-tainers with steam-sterilized SUN GRO LP5 Sunshine mix and planted each with four seeds of either A17 or DZA. These seeds had been sterilized in 10% bleach, razor-blade scarified, and cold, dark stratified on petri plates for 2 days at 4°C. After two weeks plants were thinned to ~3 plants per pot and were provided with nitrogen-free fertilizer (13) and sterile $H_2O$ as needed. These plants spent the first month of life in a growth chamber and then were moved to the greenhouse.

After twelve weeks we harvested and pooled all nodules from each replicate pot, sterilized them for 30 seconds in 10% bleach, rinsed them thoroughly, and placed the nodules in a 15 ml falcon tube with 3 ml of sterile 0.85% NaCl solution. We homogenized nodules for 1-2 min at the highest speed using a handheld tissue homogenizer (Omni International, Kennesaw, GA). We enriched for undifferentiated rhizobia (critical for obtaining measures of relative fitness) using centrifugation: first a 10 min slow spin at 400 × g to remove the heavy plant and bacteroid debris and then subjected the supernatant to a fast spin at 16,000 × g to pellet the remaining undifferentiated rhizobial cells. These pellets were frozen and we used a Plant DNeasy kit to extract DNA from both nodule pellets and the initial inoculum.

Next we generated NexteraXT libraries obtained from 250 bp paired-end reads on an Illumina NextSeq to ~120x coverage. After trimming and cleaning, SNP variant frequencies were called by aligning reads to the USDA 1106 reference genome. Sequencing revealed an unintended strain of *Ensifer medicae* in the community. This *E. medicae* strain comprised less than 0.3% of the original mixture and less than .01% of the final nodule community in either host. We omitted this strain from further analysis. We then used the Haplotype Analysis of Reads in Pools (HARP) (14) method to infer the relative frequency of each strain in each pool. This method calculates the probability that each read came from each of the 89 strains, and then determines the combination of strain frequencies that maximizes the likelihood across all reads. In other words, HARP uses all of the reads obtained from a sample to make the best estimate of the frequency of each of the 89 strains. HARP thus enables us to measure **rhizobium relative fitness**, calculated as the fold change in the frequency of a strain ($q_x$) after selection relative to its mean frequency across the four sequencing replicates of the initial community $log_2(q_x selected/q_{\bar{x}} initial)$. This transformation allows us to account for minor differences in among-strain frequencies in the initial population. For instance, strain A may have started at a frequency of .01 while strain B started at .008, and thus, this transformation levels the playing field. Additionally, this transformation ensures data are normally distributed around no change (0), thus, produces a more amenable data distribution for GWAS. Raw read data can be found at NCBI BioProject: PRJNA388336.

**Phenotypic analyses.** For each linear regression, we calculated the Pearson correlation coefficients using the "complete observation" option of the cor function in the corrplot package (v0.88, 15), and calculated adjusted p-values using the "fdr" (false discovery rate) method of the corr.test function of R's psych package (v1.8.12, 16). To visualize the resulting genetic correlations, we used a custom function that utilizes ggplot2 (v3.3.3, 17) to produce the paneled figures. Our regression analyses only included the 89 strains overlapping in both the single- and multi-strain experiments.

**DNA isolation, whole-genome sequencing, and variant calling.** In order to obtain DNA from each of the 191 rhizobia isolates, we grew overnight cultures as described above, and used the Qiagen DNeasy kit (Hilden, Germany) to extract and purify DNA, as per the manufacturer's protocol. We then submitted DNA samples to be sequenced by the DOE Joint Genome Institute (JGI) for sequencing (Berkeley, CA, USA), using the Illumina HiSeq-2500 1TB platform (Illumina, Inc., San Diego, CA, USA) with 100 bp paired-end reads. We recovered only 166 whole-genome sequences that passed quality thresholds, and thus, we regrew 25 of the 191 strains, extracted their DNA using the Zymogen Quick-DNA kit for Fungi or Bacteria (Irvine, CA, USA), and then submitted these samples to be sequenced by the Roy J. Carver Biotechnology Center at the University of Illinois at Urbana-Champaign (USA) using the Novaseq 6000 platform (Illumina, Inc, San Diego, CA, USA ) with 150 bp paired-end reads. We passed the resulting raw reads through a standard bioinformatics pipeline involving read trimming via TrimGalore! and HTStream, alignment to the reference strain *Ensifer meliloti* UDSA 1106 (assembly ID: GCA_002197065.1) using the bwa mem algorithm with default settings (v0.7.17-r1188, 18), and variant calling using FreeBayes (v1.3.1-16, 19). We hard-filtered the resulting 491,277 variants using vcftools (v0.1.17, 20), reducing the set to 36,526 variants with quality scores above 20, minor allele frequencies (MAF) of $\leq$ 5%, minimum mean depth of 20 and a maximum mean depth of 230, successful genotype calls for at least 80% of the strains, and removed any sites that differed between our samples and the reference strain, but were invariant among our samples.

**Genome-wide association tests.** Because association analyses rely on variants being statistically distinguishable, we first identified variants in strong linkage disequilibrium (LD), picking a representative variant within each LD group using the method employed in Epstein et al. (21). Briefly, we calculated the pairwise LD among all quality-filtered variants (N = 36,526), grouping variants in strong LD (95%), and then picking the variant within each LD group exhibiting the highest minor allele frequency, and then the least missing data (if ties were present). This LD grouping analysis resulted in 8,646 representative variants that went on to be tested in association analyses: 757 on the chromosome, 3,277 on the pSymA, and 4,612 on the pSymB. To run the GWAS in GEMMA, we used the -lmm 4 option and included a standardized (-gk2 option) kinship matrix in order to control for population structure among our rhizobia strains. We used the LD-filtered variants to construct k-matrices for each genomic region separately (i.e., one for each of the chromosome, pSymA, and pSymB), given that each replicon exhibits its own distinct evolutionary history (22). To conduct GWAS for rhizobium relative fitness, we repeated the LD grouping analyses and GWAS for the 89 strain subset separately, given that variants present in the larger 191 subset may not be present in the 89 subset (and vice versa, due to minor allele frequency cut-offs). All downstream analyses only included variants overlapping between the 191 and 89 strain subset.

Although GEMMA calculates p-values for each association, these are likely to be highly conservative once corrected for multiple tests, and do not take into account differences in trait distributions or LD patterns unique to each dataset (21, 23). Thus, in order to determine the significance of each association, we conducted a permutation method similar to Epstein et al. (21), whereby we randomized phenotypes with respect to genotypes, re-ran the association analyses in GEMMA using the same parameters, and repeated this process 1000 times to produce a null distribution of effect sizes for each variant in the model. We then used the findinterval method in R to determine whether our observed effect sizes exceeded the 2.5% or 97.5% intervals of the null distribution, and if so, tagged these variants as being significantly associated with our traits of interest. After running permutation tests for all 10 host-trait combinations, 2,471 ($\sim$30%) of the 8,646 LD-filtered variants were tagged as being significantly associated with one or more traits in one or more host lines: in total, only 96 of these variants were found on the chromosome, while 998 and 1,377 were found on pSymA and pSymB, respectively. We then determined whether variants were synonymous versus nonsynonymous using SnpEff (v4.3t, 24), and extracted functional information for each gene tagged by significant variants using the intersect option in bedtools (v2.29.2, 25), including the genome annotation file available from NCBI for the reference sequence *Ensifer meliloti* USDA 1106 (assembly ID: GCA_002197065.1).

**Genomic analyses.** To detect signatures of selection acting on genes associated with rhizobium fitness and symbiotic pleitropy, we calculated four test statistics ($\pi$, Tajima's D, Fu and Li's F & D) for several subsets of genes: all genes on the pSymA or pSymB plus one on the chromosome (N = 3,103; "all" genes), genes tagged by variants included in our GWAS (N = 1,726; "GWAS" genes), genes containing variants significantly associated with one or more traits (N = 1,051; "sig" genes), and our focal genes of interest, which contain at least one pleiotropic variant underlying rhizobium fitness or symbiotic pleiotropy (N = 198 total: for rhizobium fitness, N = 10 with concordant and 85 discordant effects; for symbiotic pleiotropy, 97 with concordant and 21 with discordant effects). Test statistics calculated for all 3,103 genes appear in **SI Dataset S1**.

We only included genes on the symbiosis plasmids given that only one of our focal pleiotropic genes was found on the chromosome, whereas 93 were found on pSymA and 104 on pSymB. In order to control for unequal sampling sizes within each focal gene category, we conducted t-tests comparing each focal gene category to a random subsample of equivalent size, p-values recorded, and then repeated 1000 times to calculate the proportion of tests in which our focal and randomized gene subsets were not significantly different (i.e., $P > 0.05$). If the proportion of non-significant tests was less than 10%, then we considered the focal gene category to be significantly different from the randomized set. Because we found that both "GWAS" and "sig" genes had elevated values of diversity compared to the entire genome (**Table S1**), likely because these genes by definition had to contain variants to be included in our GWAS, we decided to take a conservative approach by comparing our focal genes categories to the "sig" genes as our null.

#### SI Results

**Genomic basis of rhizobium fitness and symbiotic pleiotropy.** In total, we found 278 variants associated with pleiotropy: 110 and 148 for rhizobium fitness and symbiotic pleiotropy, respectively, while 20 overlapped both categories. These variants were found in 198 unique coding genes: 70 and 99 for rhizobium fitness and symbiotic pleiotropy, respectively, while 29 overlapped both categories. Most genes were found on pSymA (N = 93) and pSymB (N = 104), while only 1 was found on the chromosome.

Regardless of pleiotropic category (i.e., rhizobium fitness vs symbiotic), genes tagged by significant variants were often implicated in biological processes based on UNIPRO keywords that included "amino acid transport" and "transcriptional regulation", as well as molecular functions including "oxidoreductase", "transferase", "DNA binding", "lyase", and "hydrolase", suggesting that the overlap in gene networks and functions may be more common across pleiotropic categories, which could be more formally tested in follow up transcriptomic studies and gene co-expression network analyses. We discuss genes within each pleiotropic category, as well as "universal" genes spanning both, in turn below.

For loci associated with rhizobium fitness pleiotropy (N = 99 total genes; **SI Table S2 & Dataset S3**), we found several variants of interest in some the more well-known symbiosis genes on pSymA, including *noe*A (nodulation protein, ORF/RefSeq ID: SMa0773/NP_435662.1) and *fix*N (SMa1220/NP_435911.1), underlying the tradeoff between nodule number and nodule weight or rhizobium relative fitness, respectively. We found a strong signal of tradeoffs between nodule number and nodule weight in genes on pSymA involved in catalytic activity (e.g., Fumarylacetoacetate hydrolase, SMa0247/NP_435377.1), cellular membrane integrity (e.g., *dct*Q TRAP permease, SMa0249/NP_435378.1), as well as genes on pSymB involved in oxidoreductase activity (e.g., *thu*B, SM_b20330/NP_436856.1). Several genes contained multiple variants that were associated both with discordant pleiotropy between nodule number and either nodule weight or rhizobium relative fitness, as well as concordant pleiotropy between nodule weight and rhizobium relative fitness, one of which was the only gene of interest found on the chromosome (i.e., *snd*H L-sorbosone dehydrogenase, SMc04291/NP_386155.1), while others were found on pSymB (e.g., methanol dehydrogenase protein, SM_b20173/NP_436713.1; uncharacterized domain-containing protein, SM_b20139/NP_436679.1).

We found several putative symbiotic pleiotropic loci (N = 128 total genes) in canonical symbiosis genes on pSymA, including *fix* genes (I2: SMa0621/ NP_435571.1; K2: SMa0762/NP_435654.1; O3: SMa0615/NP_435567.1), *nod*H (SMa0851/NP_435710.1), and *nif*K (SMa0829/NP_435697.1), all of which underlie fitness alignment (**SI Table S2 & Dataset S3**). For example, the major allele of a variant located within the *fix*I2 gene was associated with an increase in both nodule weight and shoot biomass, while the major allele of a variant in *fix*K2 was associated with an increase in both shoot biomass and nodule number, both on DZA (**SI Dataset S2**). Genes underlying fitness conflict included a fatty acid desaturase (SMa0130/NP_435314.1), Pyrroline-5-carboxylate reductase (SM_b20003/NP_436546.1), and aldehyde dehydrogenase (SMa1844/NP_436261.1), all on A17, while a carbohydrate kinase (SM_b20489/NP_437011.1), DNA helicase (*uvr*D2: SMa2321/NP_436495.1), and esterase (SM_b21424/NP_437792.1) underlie conflict on DZA. In general, we found a strong signal in branched-chain amino acid transport and many aminotransferases for compounds important in bacteroids and symbiosis including glutamine, aspartate, asparagine, proline, and putrescine, suggesting that there is lots of potential for alignment and conflict in amino acid cycling pathways, mutants of which have been previously demonstrated to produce important symbiotic phenotypes (26).

For genes containing variants that were categorized as being "universally pleiotopic" (N = 29 total genes), most (23/29 or 79%) were located on pSymB (**SI Table S2**, **SI Dataset S3**). The major allele of these variants tended to be associated with a decrease in nodule number, and an increase in other rhizobium or plant fitness proxies (**SI Dataset S3**). Universal genes of interest included a ferrodoxin reductase, *moc*F (SM_b20820/NP_437100.1); dihydroxy-acid dehydratase, *ilv*D4 (SM_b20115/NP_436655.1); and diaminobutyric acid deacetylase, *doe*B (SM0020_02920/WP_003525897.1), all on pSymB. The only characterized protein found on pSymA was an ornithine decarboxylase (Smed_5195/YP_001313905.1). Finally, we found many pleiotropic variants in genes involved in processes that are not intuitively linked to symbiosis, such as selenocysteine synthase, molybdoterin, conjugative transfer and plasmid stabilization, hemolysin, and siderophores.

#### SI Datasets

##### SI Dataset S1 (all_gene_stats.wREADME.xlsx)

Summary table for all genes on pSymA and pSymB (as well as one on the chromosome) in which neutrality statistics were calculated (N = 3,103 genes total). The first tab "all_gene_stats" provides gene-level information, while the "README" tab provides a brief description of each column in the first tab.

##### SI Dataset S2 (pleio_vars.ps_wREADME.xlsx)

Summary table for all variants underlying rhizobium fitness (N = 130) or symbiotic (N = 168) pleiotropy. The first tab "pleio_vars.ps" provides variant-level information, while the "README" tab provides a brief description of each column in the first tab.

##### SI Dataset S3 (pleio_vars.gene_Uniprot_wREADME.xlsx)

Summary table for all genes underlying rhizobium fitness (N = 99) or symbiotic (N = 128) pleiotropy. The first tab "pleio_vars.gene" provides gene-level information, while the "README" tab provides a brief description of each column in the first tab.

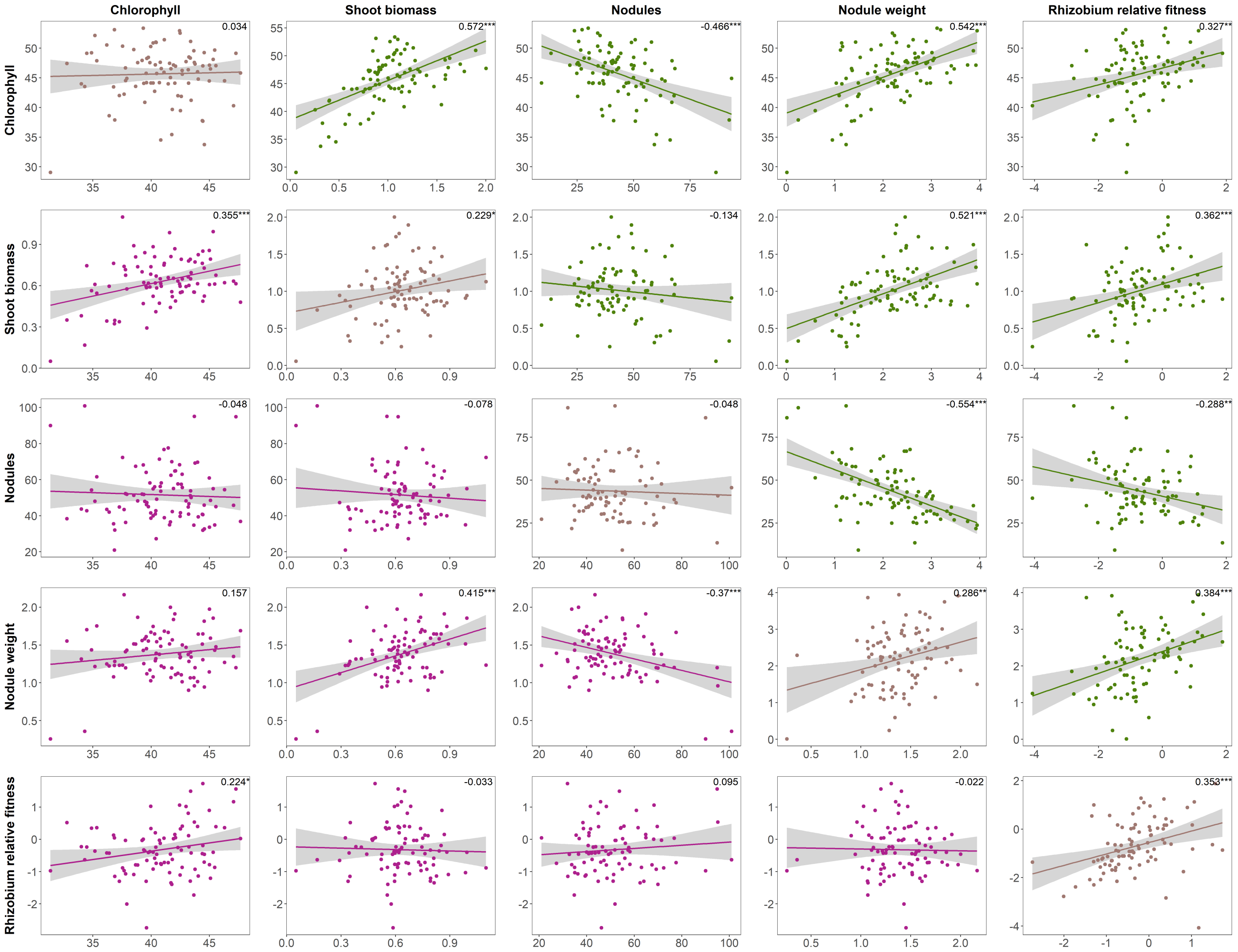


**Fig. S1. Genetic correlations for fitness proxies measured on both host plant lines.** Genetic correlations among fitness proxies measured on host lines DZA (above diagonal, in green) or A17 (below diagonal, in pink) as well as between the same fitness proxy across hosts (along diagonal). Correlations based on strain estimated marginal means corrected for rack. Numbers in top right corners of each plot indicate Pearson correlation coefficients. Significance: $p < 0.001$ = '***'; $p < 0.01$ = '**'; $p < 0.05$ = '*'.

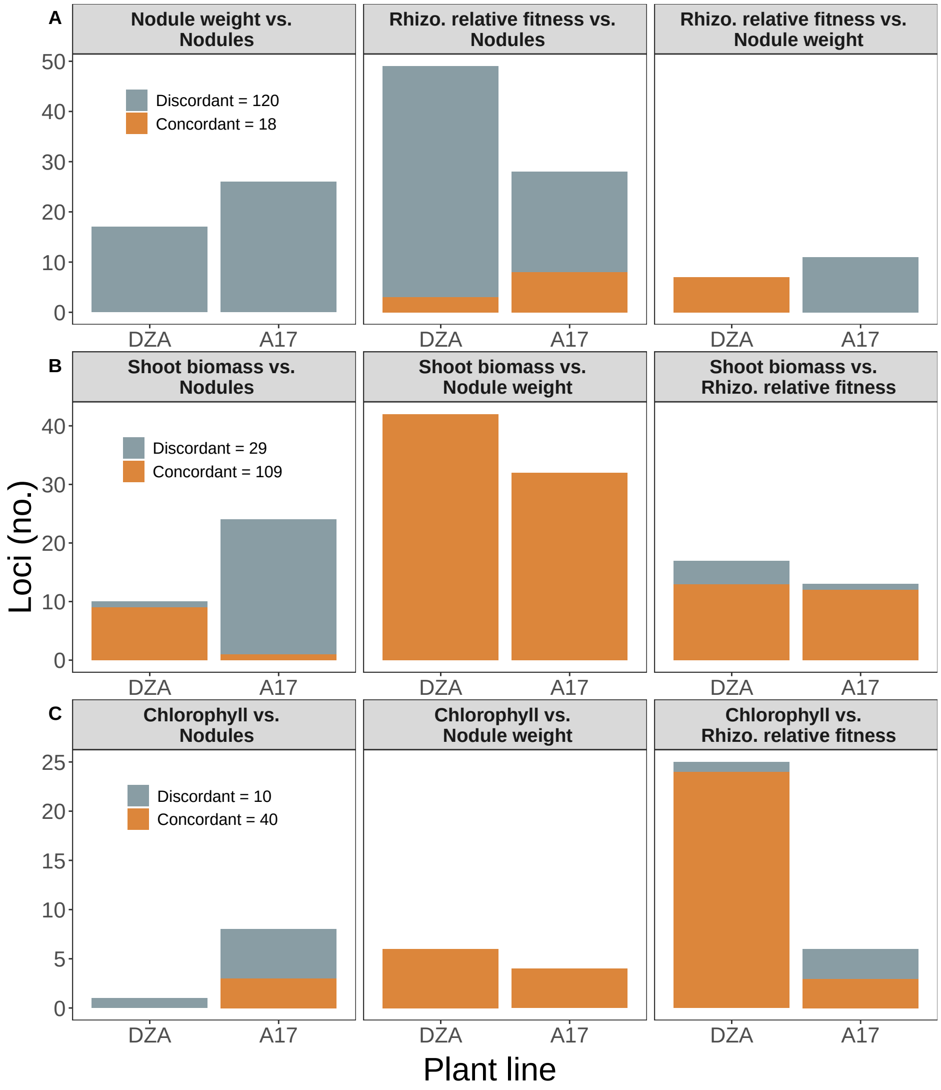


**Fig. S2. Rhizobium fitness (A) and symbiotic pleiotropy (B,C) dominated by variants with discordant (grey) or concordant (orange) effects, respectively.** Variants significantly associated with two or more rhizobium fitness proxies (nodule number, nodule size, rhizobium relative fitness) are considered to be associated with rhizobium fitness pleiotropy while those associated with both plant (shoot biomass, leaf chlorophyll A content) and rhizobium fitness proxies are associated with symbiotic pleiotropy. Total pleiotropic variant counts with same sign (i.e., concordant, orange) or opposite sign (i.e., discordant, grey) effects on both fitness proxies appear in the leftmost panels in A-C.

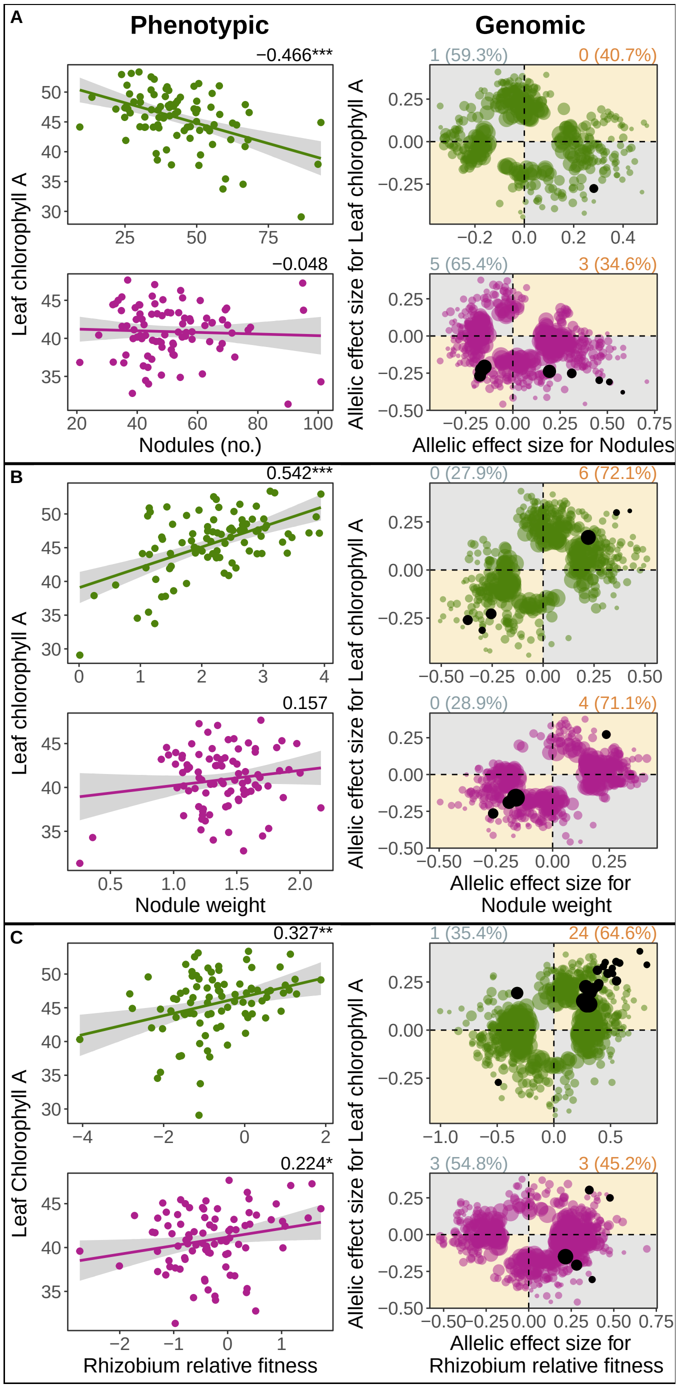


**Fig. S3. Fitness alignment between rhizobia (*Ensifer meliloti*) and host (*Medicago truncatula*) prevails at both the phenotypic (left panels) and genomic (right panels) levels. Phenotypic**: Genetic correlations between pairwise fitness proxies measured on plant lines DZA (green, top rows) or A17 (pink, bottom rows), based on 89 *Ensifer meliloti* strains. Dots represent estimated marginal strain means for leaf chlorophyll A content, as well as nodule number and nodule weight, all being measured in single-strain experiments, or medians for rhizobium relative fitness measured in multi-strain experiments. Numbers at top right of each correlation represent Pearson correlation coefficients, while asterisks represent significance: * = $P < 0.05$; ** = $P < 0.01$; *** = $P < 0.001$. **Genomic**: Dots represent allelic effect sizes (i.e., beta scores calculated in GEMMA), those falling along the diagonal (orange quadrants) or off-diagonal (grey quadrants) represent variants with concordant or discordant effects, respectively. Coloured dots represent variants that were significantly associated with one of the two fitness proxies, while black dots represent pleiotropic variants, i.e., significantly associated with both fitness proxies. Numbers outside of and percentages within parentheses at the top left and right of each plot represent the pleiotropic variant counts and the proportion of total significant variants, respectively, that are discordant (left, in grey) or concordant (right, in orange).

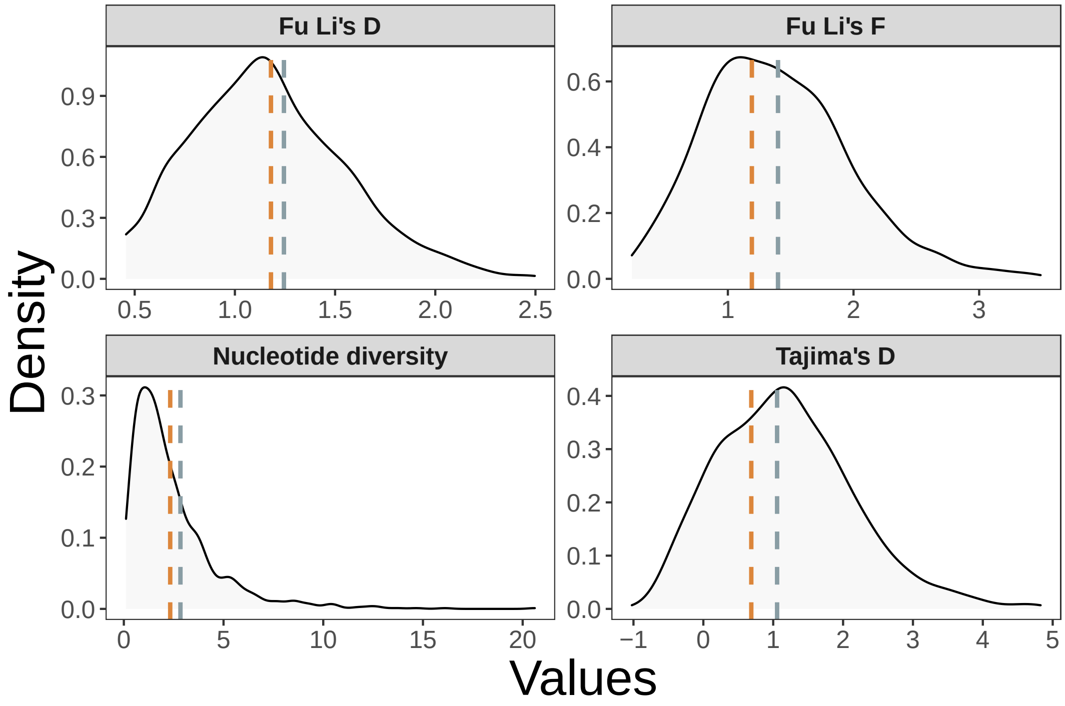


**Fig. S4. Neutrality statistics do not significantly differ from null for genes underlying rhizobium fitness pleiotropy.** Vertical lines represent the average values calculated for four separate statistics, grey for discordant and orange for concordant genes. Distributions represent the same statistics calculated for all genes containing significant variants based on GWAS. Dashed and solid lines represent non-significant ($p > 0.1$) and significant ($p < 0.1$) differences, respectively, between each focal gene category and all significant genes (i.e., distributions).

**Table S1. Neutrality statistics are elevated only for genes underlying concordant symbiotic pleiotropy.** Four different test statistics were calculated for six different gene categories: all genes on the pSymA or pSymB plus one on the chromosome (“all” genes), genes containing variants included in our GWAS (“GWAS” genes), genes containing variants significantly associated with one or more traits (“sig” genes), and our focal genes of interest, which contain at least one pleiotropic variant underlying rhizobium fitness or symbiotic pleiotropy (N = 198 genes in total). The numbers outside of parentheses for each test statistic represent the mean value, while numbers within parentheses represent P-values. Bold values are significantly different ($P < 0.10$) from the "null": for the focal gene categories (e.g., rhizobium fitness, concordant), the null is represented by the "sig" gene category, while for the GWAS and sig categories, the null is represented by the "all" gene category.

| Category | Genes (no.) | $\pi$ | Tajima’s D | Fu & Li’s D | Fu & Li’s F |
|---|---|---|---|---|---|
| all | 3,103 | 1.304 (NA) | 1.074 (NA) | 1.006 (NA) | 1.223 (NA) |
| GWAS | 1,726 | 2.069 (0) | 1.148 (0.518) | 1.068 (0.001) | 1.302 (0.004) |
| sig | 1,051 | 2.43 (0) | 1.184 (0.314) | 1.164 (0) | 1.394 (0) |
| rhizobium fitness, concordant | 10 | 2.325 (1) | 0.686 (0.767) | 1.18 (0.999) | 1.191 (0.956) |
| rhizobium fitness, discordant | 85 | 2.838 (0.905) | 1.056 (0.936) | 1.245 (0.863) | 1.399 (0.994) |
| symbiotic, concordant | 97 | **3.603 (0.041)** | 1.365 (0.768) | **1.388 (0.001)** | **1.642 (0.051)** |
| symbiotic, discordant | 21 | 2.458 (0.973) | 0.933 (0.93) | 1.258 (0.939) | 1.356 (0.992) |

**Table S2.** Gene-level summary of pleiotropic variants categorized as underlying rhizobium (rhizo.) fitness (N = 99 genes) or symbiotic (N = 128 genes) pleiotropy, and whether such pleiotropy was concordant (i.e., same-sign effects on both proxies) or discordant (i.e., opposite-sign effects on both proxies); the 'mixed' category represents genes for which some variants were concordant and others discordant. The number of variants (vars.) and associations (assocs.) represents the number of unique pleiotropic variants and the total number of associations (i.e., a single variant can have multiple associations), respectively. The 'Max effect size' represents the absolute maximum allelic effect size calculated by GEMMA of a variant contained within that gene. 'Host genotype' refers to the host in which the variant effects were significant, either on plant line DZA, A17, or both. Genes highlighted in red are mentioned in the SI Results.

| Region | RefSeq ID | Protein ID | Putative function (NCBI) | Vars. (no.) | Assocs. (no.) | Max effect size | Host genotype | Rhizo. fitness pleiotropy | Symbiotic pleiotropy |
|---|---|---|---|---|---|---|---|---|---|
| pSymA | NP_436481.1 | ASP60956.1 | penicillin-binding protein 1A | 1 | 2 | 0.263 | DZA | concordant | NA |
| pSymB | NP_436905.1 | ASP61943.1 | aspartate aminotransferase family protein | 2 | 4 | 0.330 | A17 | concordant | NA |
| pSymB | NP_437823.1 | ASP62799.1 | hypothetical protein | 1 | 2 | 0.389 | A17 | concordant | NA |
| pSymB | NP_436738.1 | ASP61779.1 | pyrroloquinoline quinone biosynthesis protein B | 1 | 3 | 0.227 | DZA | concordant | NA |
| pSymB | NP_437079.1 | ASP62112.1 | copper chaperone | 1 | 2 | 0.239 | DZA | concordant | NA |
| pSymB | NP_437156.1 | ASP62180.1 | DNA ligase D | 1 | 2 | 0.408 | DZA | concordant | NA |
| pSymB | NP_437856.1 | ASP62833.1 | cytochrome ubiquinol oxidase subunit I | 1 | 2 | 0.429 | DZA | concordant | NA |
| pSymA | NA | ASP61170.1 | hypothetical protein | 1 | 2 | 0.300 | A17 | discordant | NA |
| pSymA | NP_435272.1 | ASP60870.1 | AGE family epimerase/isomerase | 1 | 3 | 0.227 | A17 | discordant | NA |
| pSymA | NP_435292.1 | ASP60850.1 | BA14K family protein | 1 | 2 | 0.392 | A17 | discordant | NA |
| pSymA | NP_435377.1 | ASP60762.1 | fumarylacetoacetate hydrolase | 1 | 2 | 0.303 | A17 | discordant | NA |
| pSymA | NP_435378.1 | ASP60761.1 | TRAP transporter permease (*dct*Q) | 1 | 2 | 0.337 | A17 | discordant | NA |
| pSymA | NP_435547.2 | ASP60601.1 | hypothetical protein | 1 | 3 | 0.409 | A17 | discordant | NA |
| pSymA | NP_435911.1 | ASP60265.1 | cytochrome c oxidase subunit 1 (*fix*N) | 1 | 3 | 0.271 | A17 | discordant | NA |
| pSymA | NP_435919.1 | ASP60257.1 | nitrate reductase (*nap*B) | 1 | 3 | 0.239 | A17 | discordant | NA |
| pSymA | NP_436056.1 | ASP61352.1 | oxidoreductase | 1 | 2 | 0.291 | A17 | discordant | NA |
| pSymA | NP_436085.1 | ASP61324.1 | cation:proton antiporter | 1 | 2 | 0.230 | A17 | discordant | NA |
| pSymA | NP_436089.2 | ASP61320.1 | GGDEF domain-containing protein | 1 | 2 | 0.258 | A17 | discordant | NA |
| pSymA | NP_436397.2 | ASP61037.1 | hemolysin-type calcium-binding protein | 2 | 6 | 0.335 | A17 | discordant | NA |
| pSymA | NP_436518.1 | ASP60920.1 | molybdopterin dehydrogenase | 2 | 4 | 0.276 | A17 | discordant | NA |
| pSymA | NP_436519.2 | ASP61494.1 | oxidoreductase | 1 | 2 | 0.347 | A17 | discordant | NA |
| pSymA | YP_001314404.1 | ASP60576.1 | hypothetical protein | 1 | 2 | 0.273 | A17 | discordant | NA |
| pSymA | YP_001314405.1 | NA | glycerol transporter | 1 | 2 | 0.325 | A17 | discordant | NA |
| pSymA | YP_001314499.1 | ASP61343.1 | GntR family transcriptional regulator | 1 | 2 | 0.252 | A17 | discordant | NA |
| pSymA | NP_435386.1 | ASP60753.1 | alcohol dehydrogenase | 1 | 2 | 0.209 | DZA | discordant | NA |
| pSymA | NP_435662.1 | ASP60498.1 | nodulation protein (*noe*A) | 1 | 2 | 0.204 | DZA | discordant | NA |
| pSymA | NP_436112.1 | ASP61295.1 | adenylate cyclase | 1 | 3 | 0.386 | DZA | discordant | NA |
| pSymA | NP_436119.2 | ASP61291.1 | LysR family transcriptional regulator | 1 | 2 | 0.358 | DZA | discordant | NA |
| pSymA | NP_436135.2 | ASP61279.1 | nuclear transport factor 2 family protein | 1 | 2 | 0.314 | DZA | discordant | NA |
| pSymA | NP_436516.2 | ASP60922.1 | class I SAM-dependent methyltransferase | 1 | 3 | 0.510 | DZA | discordant | NA |
| pSymA | WP_014990337.1 | ASP60771.1 | hypothetical protein | 1 | 2 | 0.169 | DZA | discordant | NA |
| pSymB | NP_436594.1 | ASP63021.1 | alpha/beta hydrolase | 1 | 2 | 0.378 | A17 | discordant | NA |
| pSymB | NP_436599.1 | ASP61642.1 | SAM-dependent methyltransferase | 1 | 2 | 0.292 | A17 | discordant | NA |
| pSymB | NP_436628.1 | ASP61671.1 | protein-L-isoaspartate O-methyltransferase | 1 | 2 | 0.340 | A17 | discordant | NA |
| pSymB | NP_436773.1 | ASP61817.1 | glycosyl transferase | 1 | 2 | 0.164 | A17 | discordant | NA |
| pSymB | NP_436856.1 | ASP61896.1 | gfo/Idh/MocA family oxidoreductase (*thu*B) | 2 | 4 | 0.178 | A17 | discordant | NA |
| pSymB | NP_437315.1 | ASP62325.1 | nucleotide-binding protein | 1 | 2 | 0.247 | A17 | discordant | NA |
| pSymB | NP_437322.1 | ASP62332.1 | adenine deaminase | 1 | 2 | 0.234 | A17 | discordant | NA |
| pSymB | NP_437324.1 | ASP62334.1 | phosphoribosyltransferase | 1 | 2 | 0.221 | A17 | discordant | NA |
| pSymB | NP_437327.1 | ASP62337.1 | allantoicase | 1 | 2 | 0.260 | A17 | discordant | NA |
| pSymB | NP_437330.1 | ASP62340.1 | xanthine dehydrogenase small subunit | 1 | 2 | 0.303 | A17 | discordant | NA |
| pSymB | NP_437331.1 | ASP62341.1 | xanthine dehydrogenase molybdopterin binding subunit | 1 | 2 | 0.270 | A17 | discordant | NA |
| pSymB | NP_437386.1 | ASP62394.1 | LysR family transcriptional regulator | 2 | 4 | 0.302 | A17 | discordant | NA |
| pSymB | NP_437405.1 | ASP62411.1 | sigma-54-dependent Fis family transcriptional regulator | 1 | 2 | 0.271 | A17 | discordant | NA |
| pSymB | NP_437417.1 | ASP62422.1 | phosphoethanolamine transferase | 1 | 2 | 0.319 | A17 | discordant | NA |
| pSymB | NP_437418.1 | ASP62423.1 | hypothetical protein | 1 | 2 | 0.280 | A17 | discordant | NA |
| pSymB | YP_001312350.1 | ASP62058.1 | transporter | 1 | 2 | 0.184 | A17 | discordant | NA |
| pSymB | YP_001313347.1 | ASP62335.1 | NCS2 family permease | 1 | 2 | 0.209 | A17 | discordant | NA |
| pSymB | NP_436547.1 | ASP61595.1 | LysR family transcriptional regulator | 1 | 3 | 0.399 | DZA | discordant | NA |
| pSymB | NP_436564.1 | ASP61612.1 | Lrp/AsnC family transcriptional regulator | 1 | 2 | 0.326 | DZA | discordant | NA |
| pSymB | NP_436573.1 | ASP61621.1 | RNA polymerase sigma factor (*sig*J) | 1 | 3 | 0.263 | DZA | discordant | NA |
| pSymB | NP_436656.1 | ASP61697.1 | amylo-alpha-1,6-glucosidase | 2 | 6 | 0.412 | DZA | discordant | NA |
| pSymB | NP_436680.1 | ASP61722.1 | dihydrodipicolinate synthase family protein | 1 | 3 | 0.454 | DZA | discordant | NA |
| pSymB | NP_436709.1 | ASP61751.1 | hypothetical protein | 1 | 2 | 0.323 | DZA | discordant | NA |
| pSymB | NP_436710.1 | ASP61752.1 | S-(hydroxymethyl)glutathione dehydrogenase | 1 | 2 | 0.331 | DZA | discordant | NA |
| pSymB | NP_436766.1 | ASP61809.1 | ABC transporter ATP-binding protein | 1 | 2 | 0.448 | DZA | discordant | NA |
| pSymB | NP_436789.1 | ASP61832.1 | dihydrodipicolinate synthase family protein | 1 | 2 | 0.209 | DZA | discordant | NA |
| pSymB | NP_436791.1 | ASP61834.1 | malate dehydrogenase | 1 | 2 | 0.267 | DZA | discordant | NA |
| pSymB | NP_436797.1 | ASP61840.1 | amino acid dehydrogenase | 1 | 2 | 0.343 | DZA | discordant | NA |
| pSymB | NP_436798.1 | ASP61841.1 | hydroxyproline-2-epimerase | 1 | 2 | 0.378 | DZA | discordant | NA |
| pSymB | NP_437053.1 | ASP62083.1 | sugar phosphate isomerase/epimerase | 1 | 3 | 0.358 | DZA | discordant | NA |
| pSymB | NP_437093.2 | ASP62125.1 | ABC transporter ATP-binding protein | 1 | 3 | 0.256 | DZA | discordant | NA |

Table S2 continued from previous page

| Region | RefSeq ID | Protein ID | Putative function (NCBI) | Vars. (no.) | Assocs. (no.) | Max effect size | Host genotype | Rhizo. fitness pleiotropy | Symbiotic pleiotropy |
|---|---|---|---|---|---|---|---|---|---|
| pSymB | NP_437094.1 | ASP62126.1 | hypothetical protein | 1 | 2 | 0.397 | DZA | discordant | NA |
| pSymB | NP_438092.1 | ASP61570.1 | Putative enoyl-CoA hydratase protein (*paa*G) | 1 | 2 | 0.250 | DZA | discordant | NA |
| pSymB | YP_001312688.1 | ASP61753.1 | S-formylglutathione hydrolase | 1 | 2 | 0.238 | DZA | discordant | NA |
| pSymB | YP_002122330.1 | ASP62043.1 | DUF899 domain-containing protein | 1 | 2 | 0.519 | DZA | discordant | NA |
| pSymB | NP_436767.1 | ASP61810.1 | hypothetical protein | 2 | 5 | 0.489 | Both | discordant | NA |
| Chrom | NP_386155.1 | ASP59485.1 | sorbosone dehydrogenase (*snd*H) | 1 | 3 | 0.423 | DZA | mixed | NA |
| pSymB | NP_436679.1 | ASP61721.1 | DUF2243 domain-containing protein | 2 | 6 | 0.286 | DZA | mixed | NA |
| pSymB | NP_436713.1 | ASP61755.1 | PQQ-dependent dehydrogenase, methanol/ethanol family | 1 | 4 | 0.423 | DZA | mixed | NA |
| pSymA | NP_435247.1 | ASP60899.1 | formate dehydrogenase-N subunit alpha | 1 | 2 | 0.257 | A17 | NA | concordant |
| pSymA | NP_435250.1 | ASP60896.1 | formate dehydrogenase accessory protein (*fdh*E) | 4 | 8 | 0.279 | A17 | NA | concordant |
| pSymA | NP_435251.1 | ASP60895.1 | L-selenocysteinyl-tRNA(Sec) synthase | 3 | 6 | 0.291 | A17 | NA | concordant |
| pSymA | NP_435253.1 | ASP60893.1 | selenocysteine-specific translation elongation factor | 1 | 2 | 0.356 | A17 | NA | concordant |
| pSymA | NP_435307.1 | ASP60834.1 | cytochrome C | 1 | 2 | 0.214 | A17 | NA | concordant |
| pSymA | NP_435593.1 | ASP60560.1 | hypothetical protein | 2 | 4 | 0.327 | A17 | NA | concordant |
| pSymA | NP_435594.1 | ASP60559.1 | transcriptional regulator | 1 | 2 | 0.298 | A17 | NA | concordant |
| pSymA | NP_435599.1 | ASP60555.1 | HlyD family secretion protein | 1 | 3 | 0.166 | A17 | NA | concordant |
| pSymA | NP_435610.2 | ASP60545.1 | hypothetical protein | 1 | 2 | 0.191 | A17 | NA | concordant |
| pSymA | NP_435697.1 | ASP60464.1 | nitrogenase molybdenum-iron protein subunit beta (*nif*K) | 1 | 3 | 0.270 | A17 | NA | concordant |
| pSymA | NP_435710.1 | ASP60452.1 | nodulation protein H (*nod*H) | 1 | 2 | 0.186 | A17 | NA | concordant |
| pSymA | NP_436262.1 | ASP61163.1 | tartrate dehydrogenase | 1 | 2 | 0.195 | A17 | NA | concordant |
| pSymA | NP_436462.1 | ASP60971.1 | integrase | 2 | 4 | 0.333 | A17 | NA | concordant |
| pSymA | NP_435259.1 | ASP60883.1 | hypothetical protein | 1 | 2 | 0.244 | DZA | NA | concordant |
| pSymA | NP_435294.1 | ASP60848.1 | D-aminopeptidase | 1 | 2 | 0.256 | DZA | NA | concordant |
| pSymA | NP_435306.1 | ASP60835.1 | hypothetical protein | 1 | 2 | 0.328 | DZA | NA | concordant |
| pSymA | NP_435317.2 | ASP60823.1 | AI-2E family transporter | 1 | 2 | 0.246 | DZA | NA | concordant |
| pSymA | NP_435318.2 | ASP60822.1 | bifunctional diguanylate cyclase/phosphodiesterase | 1 | 2 | 0.259 | DZA | NA | concordant |
| pSymA | NP_435324.1 | ASP60817.1 | hypothetical protein | 1 | 2 | 0.276 | DZA | NA | concordant |
| pSymA | NP_435326.1 | ASP60815.1 | malonyl-CoA synthase | 1 | 3 | 0.204 | DZA | NA | concordant |
| pSymA | NP_435334.1 | ASP60807.1 | hypothetical protein | 1 | 2 | 0.168 | DZA | NA | concordant |
| pSymA | NP_435343.1 | ASP60798.1 | MFS transporter | 1 | 2 | 0.185 | DZA | NA | concordant |
| pSymA | NP_435368.1 | ASP60772.1 | glutamate dehydrogenase | 1 | 2 | 0.237 | DZA | NA | concordant |
| pSymA | NP_435372.1 | ASP60767.1 | dihydroxy-acid dehydratase | 1 | 2 | 0.259 | DZA | NA | concordant |
| pSymA | NP_435559.2 | ASP60593.1 | alpha/beta hydrolase | 1 | 2 | 0.281 | DZA | NA | concordant |
| pSymA | NP_435560.2 | ASP60592.1 | adenylate cyclase | 1 | 2 | 0.299 | DZA | NA | concordant |
| pSymA | NP_435567.1 | ASP60586.1 | cytochrome c oxidase cbb3-type subunit II (*fix*O3) | 1 | 3 | 0.185 | DZA | NA | concordant |
| pSymA | NP_435571.1 | ASP60581.1 | copper-translocating P-type ATPase (*fix*I2) | 1 | 2 | 0.237 | DZA | NA | concordant |
| pSymA | NP_435604.2 | ASP60550.1 | amino acid ABC transporter substrate-binding protein | 1 | 3 | 0.196 | DZA | NA | concordant |
| pSymA | NP_435615.1 | ASP60540.1 | carbamate kinase | 1 | 2 | 0.211 | DZA | NA | concordant |
| pSymA | NP_435621.1 | ASP60534.1 | sugar ABC transporter substrate-binding protein | 2 | 4 | 0.388 | DZA | NA | concordant |
| pSymA | NP_435623.1 | ASP60532.1 | carbohydrate ABC transporter permease | 2 | 4 | 0.224 | DZA | NA | concordant |
| pSymA | NP_435628.1 | ASP61474.1 | NAD(P)-dependent oxidoreductase | 1 | 2 | 0.308 | DZA | NA | concordant |
| pSymA | NP_435641.1 | ASP60517.1 | molecular chaperone (*gro*EL) | 1 | 2 | 0.189 | DZA | NA | concordant |
| pSymA | NP_435654.1 | ASP60507.1 | transcriptional regulator (*fix*K2) | 1 | 2 | 0.171 | DZA | NA | concordant |
| pSymA | NP_435748.1 | ASP60415.1 | Ti-type conjugative transfer system protein (*tra*G) | 1 | 2 | 0.206 | DZA | NA | concordant |
| pSymA | NP_435752.1 | ASP60411.1 | mechanosensitive ion channel family protein | 1 | 2 | 0.452 | DZA | NA | concordant |
| pSymA | NP_435753.1 | ASP60410.1 | sensor histidine kinase | 2 | 4 | 0.384 | DZA | NA | concordant |
| pSymA | NP_435978.2 | ASP61431.1 | sugar ABC transporter permease | 1 | 2 | 0.464 | DZA | NA | concordant |
| pSymA | NP_435981.1 | ASP61427.1 | ABC transporter ATP-binding protein | 1 | 3 | 0.291 | DZA | NA | concordant |
| pSymA | NP_435985.1 | ASP61423.1 | sugar phosphate isomerase/epimerase | 1 | 2 | 0.404 | DZA | NA | concordant |
| pSymA | NP_436006.1 | ASP61401.1 | LysR family transcriptional regulator | 1 | 2 | 0.403 | DZA | NA | concordant |
| pSymA | NP_436072.1 | ASP61336.1 | NADH-quinone oxidoreductase subunit I 2 | 2 | 5 | 0.314 | DZA | NA | concordant |
| pSymA | NP_436073.1 | ASP61335.1 | histidine kinase | 1 | 2 | 0.203 | DZA | NA | concordant |
| pSymA | NP_436166.2 | ASP61523.1 | amidohydrolase family protein | 1 | 3 | 0.319 | DZA | NA | concordant |
| pSymA | NP_436209.1 | ASP61209.1 | TonB-dependent siderophore receptor | 1 | 2 | 0.276 | DZA | NA | concordant |
| pSymA | NP_436219.1 | ASP61201.1 | aminotransferase | 1 | 2 | 0.384 | DZA | NA | concordant |
| pSymA | NP_436244.1 | ASP61179.1 | MBL fold metallo-hydrolase | 1 | 2 | 0.197 | DZA | NA | concordant |
| pSymA | NP_436245.1 | ASP61178.1 | MFS transporter | 1 | 2 | 0.201 | DZA | NA | concordant |
| pSymA | NP_436271.1 | ASP61155.1 | ABC transporter permease | 1 | 3 | 0.334 | DZA | NA | concordant |
| pSymA | NP_436482.1 | ASP60955.1 | penicillin-binding protein 2 | 1 | 3 | 0.350 | DZA | NA | concordant |
| pSymA | YP_001312338.1 | ASP60533.1 | sugar ABC transporter permease | 1 | 2 | 0.206 | DZA | NA | concordant |
| pSymA | YP_001313599.1 | NA | cell envelope protein | 3 | 7 | 0.266 | DZA | NA | concordant |
| pSymA | YP_001314274.1 | ASP61511.1 | N-acetyltransferase | 2 | 6 | 0.345 | DZA | NA | concordant |
| pSymA | YP_001314472.1 | ASP60871.1 | mandelate racemase | 1 | 2 | 0.245 | DZA | NA | concordant |
| pSymA | NP_435603.2 | ASP60551.1 | calcium/proton exchanger | 3 | 11 | 0.210 | Both | NA | concordant |
| pSymB | NP_436553.1 | ASP61601.1 | alpha/beta hydrolase | 1 | 2 | 0.296 | A17 | NA | concordant |
| pSymB | NP_436569.1 | ASP61617.1 | tripartite tricarboxylate transporter TctB family protein | 1 | 3 | 0.199 | A17 | NA | concordant |
| pSymB | NP_436640.1 | ASP61682.1 | LLM class F420-dependent oxidoreductase | 1 | 3 | 0.237 | A17 | NA | concordant |
| pSymB | NP_436646.1 | ASP61688.1 | FadR family transcriptional regulator | 1 | 3 | 0.304 | A17 | NA | concordant |

Table S2 continued from previous page

| Region | RefSeq ID | Protein ID | Putative function (NCBI) | Vars. (no.) | Assocs. (no.) | Max effect size | Host genotype | Rhizo. fitness pleiotropy | Symbiotic pleiotropy |
|---|---|---|---|---|---|---|---|---|---|
| pSymB | NP_436678.1 | ASP61720.1 | amidohydrolase | 1 | 2 | 0.271 | A17 | NA | concordant |
| pSymB | NP_436774.1 | ASP61818.1 | CDP-paratose 2-epimerase | 1 | 2 | 0.165 | A17 | NA | concordant |
| pSymB | NP_436981.1 | ASP62012.1 | UDP-glucose 4-epimerase (*gal*E) | 1 | 3 | 0.276 | A17 | NA | concordant |
| pSymB | NP_437117.1 | ASP62145.1 | cell surface protein | 1 | 2 | 0.254 | A17 | NA | concordant |
| pSymB | NP_437160.1 | ASP62190.1 | hypothetical protein | 1 | 2 | 0.190 | A17 | NA | concordant |
| pSymB | NP_437221.1 | ASP62245.1 | IclR family transcriptional regulator | 1 | 4 | 0.163 | A17 | NA | concordant |
| pSymB | NP_437366.1 | ASP62374.1 | beta-N-acetylhexosaminidase | 1 | 2 | 0.364 | A17 | NA | concordant |
| pSymB | NP_437398.1 | ASP62405.1 | fructose-bisphosphate aldolase class II | 1 | 2 | 0.331 | A17 | NA | concordant |
| pSymB | NP_437637.1 | ASP62628.1 | ABC transporter ATP-binding protein | 1 | 2 | 0.195 | A17 | NA | concordant |
| pSymB | NP_437662.1 | ASP62652.1 | hypothetical protein | 1 | 3 | 0.213 | A17 | NA | concordant |
| pSymB | YP_001313261.1 | ASP62257.1 | two-component system response regulator | 1 | 2 | 0.173 | A17 | NA | concordant |
| pSymB | NP_436559.1 | ASP61606.1 | ABC transporter permease | 1 | 3 | 0.284 | DZA | NA | concordant |
| pSymB | NP_436799.1 | ASP61842.1 | hypothetical protein | 1 | 2 | 0.209 | DZA | NA | concordant |
| pSymB | NP_436969.1 | ASP62001.1 | GGDEF-domain containing protein | 1 | 2 | 0.384 | DZA | NA | concordant |
| pSymB | NP_437002.1 | ASP62033.1 | FadR family transcriptional regulator | 2 | 6 | 0.215 | DZA | NA | concordant |
| pSymB | NP_437003.1 | ASP62034.1 | N-acetylglutaminylglutamine amidotransferase | 1 | 2 | 0.193 | DZA | NA | concordant |
| pSymB | NP_437007.1 | ASP62038.1 | sugar ABC transporter ATP-binding protein | 1 | 3 | 0.214 | DZA | NA | concordant |
| pSymB | NP_437015.1 | ASP62046.1 | 3-ketoacyl-ACP reductase | 1 | 3 | 0.290 | DZA | NA | concordant |
| pSymB | NP_437016.1 | ASP62047.1 | LysR family transcriptional regulator | 1 | 3 | 0.228 | DZA | NA | concordant |
| pSymB | NP_437030.1 | ASP62061.1 | arabinose ABC transporter substrate-binding protein | 1 | 2 | 0.197 | DZA | NA | concordant |
| pSymB | NP_437035.1 | ASP62067.1 | chemotaxis protein (*che*B) | 1 | 2 | 0.262 | DZA | NA | concordant |
| pSymB | NP_437145.1 | ASP62168.1 | hypothetical protein | 1 | 2 | 0.290 | DZA | NA | concordant |
| pSymB | NP_437243.1 | ASP62266.1 | acetyl/propionyl/methylcrotonyl-CoA | 1 | 2 | 0.249 | DZA | NA | concordant |
| pSymB | NP_437510.1 | ASP62509.1 | sulfoacetaldehyde acetyltransferase | 1 | 3 | 0.355 | DZA | NA | concordant |
| pSymB | NP_437900.1 | ASP62873.1 | asparagine synthetase B | 1 | 2 | 0.407 | DZA | NA | concordant |
| pSymA | NP_435314.1 | ASP60827.1 | fatty acid desaturase | 1 | 2 | 0.176 | A17 | NA | discordant |
| pSymA | NP_435598.1 | ASP60556.1 | hypothetical protein | 1 | 2 | 0.187 | A17 | NA | discordant |
| pSymA | NP_436261.1 | ASP61164.1 | aldehyde dehydrogenase, NAD(P)-dependent | 1 | 2 | 0.281 | A17 | NA | discordant |
| pSymA | NP_436495.1 | ASP60944.1 | ATP-dependent helicase (*uvr*D2) | 1 | 2 | 0.279 | DZA | NA | discordant |
| pSymB | NP_436546.1 | ASP61594.1 | pyrroline-5-carboxylate reductase | 1 | 2 | 0.161 | A17 | NA | discordant |
| pSymB | NP_436984.1 | ASP62015.1 | beta-mannosidase | 1 | 2 | 0.245 | A17 | NA | discordant |
| pSymB | NP_438036.2 | ASP63118.1 | sulfite oxidase-like oxidoreductase | 1 | 3 | 0.321 | A17 | NA | discordant |
| pSymB | NP_437005.1 | ASP62036.1 | transcriptional regulator | 1 | 2 | 0.232 | DZA | NA | discordant |
| pSymB | NP_437011.1 | ASP62041.1 | carbohydrate kinase | 1 | 2 | 0.258 | DZA | NA | discordant |
| pSymB | NP_437792.1 | ASP62771.1 | acetyl esterase | 1 | 2 | 0.302 | DZA | NA | discordant |
| pSymA | NP_435605.1 | ASP60549.1 | putrescine-ornithine antiporter | 2 | 4 | 0.236 | DZA | NA | mixed |
| pSymA | NP_435751.1 | ASP60412.1 | Ti-type conjugative transfer relaxase (*tra*A) | 2 | 5 | 0.284 | DZA | NA | mixed |
| pSymB | NP_436648.1 | ASP61689.1 | peptide ABC transporter substrate-binding protein | 2 | 6 | 0.309 | A17 | NA | mixed |
| pSymB | YP_001313723.1 | ASP62127.1 | membrane assembly protein (*asm*A) | 2 | 4 | 0.335 | Both | NA | mixed |
| pSymB | NP_436880.1 | ASP61921.1 | hypothetical protein | 2 | 5 | 0.308 | A17 | concordant | discordant |
| pSymB | NP_436558.1 | ASP61605.1 | ABC transporter permease | 1 | 4 | 0.373 | A17 | discordant | mixed |
| pSymB | NP_436561.1 | ASP61609.1 | sugar ABC transporter substrate-binding protein | 1 | 3 | 0.373 | A17 | discordant | mixed |
| pSymB | NP_436633.1 | ASP61675.1 | lysylphosphatidylglycerol synthetase family protein | 2 | 6 | 0.361 | A17 | discordant | mixed |
| pSymB | NP_436643.1 | ASP61685.1 | FAD-dependent oxidoreductase | 1 | 4 | 0.399 | A17 | discordant | mixed |
| pSymB | NP_437001.1 | ASP62032.1 | ABC transporter ATP-binding protein | 1 | 3 | 0.345 | A17 | discordant | mixed |
| pSymA | NP_436524.1 | ASP60914.1 | amidase | 2 | 5 | 0.288 | DZA | concordant | concordant |
| pSymA | NP_435596.1 | ASP60557.1 | hypothetical protein | 3 | 6 | 0.255 | DZA | discordant | concordant |
| pSymA | NP_435597.2 | ASP61477.1 | AI-2E family transporter | 4 | 9 | 0.208 | DZA | discordant | concordant |
| pSymB | NP_437804.1 | ASP62780.1 | Zn-dependent hydrolase | 2 | 5 | 0.381 | Both | concordant | discordant |
| pSymA | NP_435627.1 | ASP60529.1 | IclR family transcriptional regulator | 2 | 6 | 0.208 | Both | discordant | concordant |
| pSymA | NP_435563.1 | ASP60589.1 | hypothetical protein | 2 | 4 | 0.306 | Both | discordant | concordant |
| pSymA | YP_001313905.1 | ASP60548.1 | ornithine decarboxylase | 3 | 7 | 0.385 | Both | discordant | concordant |
| pSymB | NP_436566.1 | ASP61615.1 | 5-oxoprolinase | 2 | 4 | 0.444 | Both | discordant | concordant |
| pSymB | NP_437100.1 | ASP62131.1 | ferredoxin reductase (*moc*F) | 2 | 4 | 0.275 | Both | discordant | concordant |
| pSymB | NP_437305.1 | ASP62318.1 | peptide ABC transporter substrate-binding protein | 2 | 4 | 0.329 | Both | discordant | concordant |
| pSymB | NP_436655.1 | ASP61696.1 | dihydroxy-acid dehydratase (*ilv*D4) | 2 | 8 | 0.339 | Both | discordant | concordant |
| pSymB | NP_437004.1 | ASP62035.1 | N-acetylglutaminylglutamine synthetase | 4 | 10 | 0.470 | Both | discordant | concordant |
| pSymB | NP_436652.1 | ASP61693.1 | EamA/RhaT family transporter | 1 | 4 | 0.301 | Both | discordant | discordant |
| pSymB | NP_436701.1 | ASP61743.1 | hypothetical protein | 1 | 4 | 0.349 | Both | discordant | discordant |
| pSymB | NP_436950.1 | ASP61984.1 | NAD-dependent succinate-semialdehyde dehydrogenase | 1 | 4 | 0.559 | Both | discordant | discordant |
| pSymB | NP_436967.1 | ASP61999.1 | L-idonate 5-dehydrogenase | 1 | 4 | 0.370 | Both | discordant | discordant |
| pSymB | NP_436974.1 | ASP63043.1 | hypothetical protein | 1 | 4 | 0.452 | Both | discordant | discordant |
| pSymB | NP_436975.1 | ASP62006.1 | gluconolactonase | 2 | 9 | 0.423 | Both | discordant | discordant |
| pSymB | WP_003525897.1 | ASP61994.1 | N-alpha-acetyl diaminobutyric acid deacetylase (*doe*B) | 1 | 4 | 0.636 | Both | discordant | discordant |
| pSymB | NP_436651.1 | ASP61692.1 | ABC transporter ATP-binding protein | 2 | 7 | 0.307 | Both | discordant | discordant |
| pSymB | NP_437017.1 | ASP63047.1 | hypothetical protein | 2 | 6 | 0.566 | Both | discordant | discordant |
| pSymB | NP_436650.1 | ASP61691.1 | ABC transporter permease | 2 | 9 | 0.358 | Both | discordant | mixed |
| pSymB | NP_437206.1 | ASP62230.1 | Ti-type conjugative transfer relaxase (*tra*A) | 4 | 9 | 0.584 | Both | mixed | concordant |
